# The Urinary Tract commensal *Peptoniphilus* spp. Encodes a Novel 17β-Hydroxysteroid Dehydrogenase

**DOI:** 10.64898/2026.03.29.714803

**Authors:** Briawna Binion, Saeed Ahmad, Taojun Wang, Elizabeth Tang, Betsy Barnick, Damie Olukoya, Pauline Mbuvi, Debapriya Dutta, John W. Erdman, H. Rex Gaskins, Glen Yang, Joseph Irudayaraj, Jason M. Ridlon

## Abstract

Microbial steroid metabolism represents an underappreciated extension of the vertebrate endocrine system, with growing evidence that host-associated microbes contribute to the diversity and bioavailability of sex steroids within human tissues. Emerging studies have linked microbial androgen metabolism to urinary microbiome composition and to resistance to androgen deprivation therapy (ADT) in prostate cancer. While microbial pathways capable of converting steroid precursors such as cortisol to androgens, via the steroid-17,20-desmolase pathway, such as DesG-mediated interconversion of androstenedione to testosterone have been reported, the diversity of enzymes mediating downstream androgen interconversion remains incompletely defined. Here, we investigate the androgen-forming capabilities of anaerobic bacteria from the male genitourinary microbiome, focusing on NADPH-dependent 17β-hydroxysteroid dehydrogenases (17β-HSDHs) that catalyze interconversion of androstenedione and testosterone. We isolated androgen-forming bacterial strains from human male urine and identified a previously uncharacterized 17β-HSDH encoded by *Peptoniphilus obesi*, demonstrated that this enzyme catalyzes the NADPH-dependent reduction of androstenedione to testosterone and the reverse oxidation reaction. Sequence similarity searches further identified a homologous 17β-HSDH in *Anaerococcus*, which was synthesized and functionally validated, revealing conserved activity despite low sequence identity to the previously characterized urinary tract enzyme DesG. The enzymes were found to have broad substrate specificity for C19 and C18 17keto− and 17β-hydroxysteroids. Together, these findings expand the known diversity of microbial 17β-HSDHs and identify previously unrecognized androgen-forming activities within the genitourinary microbiome.

**Importance:** Microbial steroid-transforming pathways may provide a mechanism by which commensal anaerobes contribute to androgen availability in the genitourinary tract. By identifying novel 17β-hydroxysteroid dehydrogenases from *Peptoniphilus* and *Anaerococcus*, genera repeatedly associated with prostate cancer, this study provides mechanistic insight into how microbial steroid metabolism may influence hormone-driven disease.

## Introduction

Microbial steroid metabolism has emerged as a critical, yet understudied extension of the vertebrate endocrine system (1), expanding the diversity and activity of steroids within human-associated niches. The study of the bidirectional relationship between sex-dependent steroid metabolomes in host tissues, the immune system, and the microbiome has been termed the “microgenderome” (2). Correlations between circulating testosterone and estradiol metabolites and gut microbial taxonomic diversity and relative abundance have been reported (3, 4). Alterations in hormone receptors, such as the estrogen receptor (ER-*β*), have been shown to affect the composition of the gut microbiota (5). Sex differences in microbiome composition in mice are altered after gonadectomy and/or hormone treatment (6, 7). Fecal microbiota transplant (FMT) from male to female rodents is reported to elevate circulating testosterone, which was prevented by treatment with the androgen-receptor (AR) antagonist, flutamide (8). This observation is particularly interesting since flutamide is used to treat prostate cancer, and historically, the source of AR agonists has been assumed to be solely derived from host tissue metabolism.

Prostate cancer is a hormone-sensitive cancer whose primary treatment is androgen-deprivation therapy (ADT). Evidence for the involvement of the gut microbiome in resistance to ADT comes from a study showing that FMT from human donors with metastatic castration-resistant prostate cancer (mCRPC) promoted castration-resistance in mouse models of mCRPC (9). In that study, it was reported that bacteria can convert pregnanolone to progesterone, dehydroepiandrosterone (DHEA), androstenedione (AD), and testosterone (T). In the case of prostate cancer, conversion of non-androgens (i.e pregnanolone) to androgens (i.e., T) may promote disease progression. The genes encoding enzymes in this pathway were not reported in that study (9), but determining such enzymatic pathways could establish novel therapeutic interventions.

The gut microbiome is also capable of generating androgens from the “cortisol side-chain cleavage” pathway (10, 11). This reaction was first reported in the late 1950’s (12, 13), and gut bacteria expressing the enzyme steroid-17,20-desmolase were isolated in the 1980’s (14, 15). The genes encoding steroid-17,20-desmolase (*desAB*) and steroid 20ɑ-hydroxysteroid dehydrogenase (*desC*) in the gut bacterium *Clostridium scindens* ATCC 35704 were reported in 2013 (10). *Butyricicoccus desmolans* and *Clostridium cadavaris* were previously reported to express both steroid-17,20-desmolase (*desAB*) and steroid 20β-hydroxysteroid dehydrogenase. (16) Phylogenetic and sequence similarity network analysis of the DesA and DesB sequences identified *desAB* genes in *B. desmolans* and *C. cadavaris* along with genes encoding putative 20β-hydroxysteroid dehydrogenase, which were also found in intestinal bifidobacteria (17). Indeed, we characterized the structure and function of a short-chain dehydrogenase/reductase protein family member adjacent to *desAB* genes with 20β-hydroxysteroid dehydrogenase activity encoded by the *desE* gene (17). Phylogenetic and sequence similarity network analysis of steroid-17,20-desmolase proteins revealed that the urinary tract commensal, *Propionimicrobium lymphophilum*, encoded *desABE,* which we confirmed functionally (17, 18).

Research in recent years has both confirmed that the urinary tract harbors a commensal microbiota (19) and that the composition of the urinary microbiota differs between healthy males and males diagnosed with prostate cancer (20, 21). Whether the differential taxonomic abundances observed are a cause or consequence of the development and progression of prostate cancer is currently unclear; however, the discovery that urinary tract bacteria are capable of generating androgen precursors from urinary glucocorticoids prompted us to collect urine from males newly diagnosed with prostate cancer along with age-matched healthy control males (18, 22). Several strains of *P. lymphophilum* from both urine as well as prostate cancer tissue capable of the side-chain cleavage of cortisol and prednisone, but also conversion of these metabolites to testosterone derivatives (18). We identified and characterized the recently named *desG* gene encoding an NADPH-dependent 17β-hydroxysteroid dehydrogenase (17β-HSDH) (18, 23). Most recently, we reported identification of genes encoding a novel DHEA isomerase/reductase (*dirAB*) pathway involved in conversion of DHEA to T by the urinary tract isolate *Actinobaculum massiliense* (22).

Following these discoveries, we sought to identify additional microbial enzymes involved in androgen formation in the genitourinary microbiome. Here, we report the isolation of additional androgen forming microbial genera from human male urine. We identified and biochemically characterized two novel 17β-HSDHs, one from *Peptoniphilus* sp. and a sequence homolog in *Anaerococcus* sp. Marseille-P9784, both distinct from DesG in sequence and substrate specificity. Notably, in microbiome studies comparing urinary abundances of taxa between control subjects and patients diagnosed with prostate cancer, both *Peptoniphilus* and *Anaerococcus* are consistently associated with prostate cancer (20, 24–26) but have not previously been reported to metabolize steroids. Enzyme bioconversion assays demonstrated that each enzyme catalyzes the NADPH-dependent interconversion of androstenedione and testosterone, confirming their roles in microbial androgen metabolism. Together, these findings expand the known diversity of microbial 17β-HSDHs within host-associated niches and support a potential mechanistic link between microbial steroid transformation and hormone-sensitive urogenital diseases.

## Materials and Methods

### Bacteria and chemicals

*E. coli* BL21-CodonPlus (DE3)-RIL Competent Cells were purchased from Agilent (Santa Clara, CA, USA). The urinary tract bacterial strains *Peptoniphilus obesi* CFH08 and *Peptoniphilus harei* CFH36 were isolated in this study.

Chemicals included: cortisol (Sigma); 11-deoxycortisol (Steraloids, Newport, RI, USA); 11β-hydroxyandrostenedione (11OHAD) (Steraloids); 11-ketoandrostenedione (11KAD) (Steraloids); 11-ketotestosterone (11KT) (Steraloids); δ¹-Adrenosterone (1,4-11KAD) (Steraloids); 11β-hydroxtestosterone (11OHT) (Steraloids); Dehydroepiandrosterone (DHEA) (Steraloids); Androstenedione (AD) (Steraloids); Androstenediol (5-AL) (Steraloids); Testosterone (T) (Steraloids), 5α-androstenedione (5α-AD) (Steraloids); dihydrotestosterone (DHT) (Steraloids).

### Bacterial media preparation

Lysogeny broth (LB, Sigma) was purchased and prepared according to the manufacturer’s instructions. Peptone Yeast Glucose (PYG) broth (modified) was prepared according to the DSMZ protocol. Blood agar base (Sigma) was purchased and prepared according to the instructions, with 6% (v/v) defibrinated sheep blood (Thermo Scientific) added. Schaedler agar (Sigma) was purchased and prepared according to the manufacturer’s instructions. Columbia broth (BBL) was purchased and prepared as instructed, with 15 g/L agar added to solidify the medium.

### Patient recruitment and urine sample collection

To identify androgen forming urinary microbial taxa, we consented and recruited patients under IRB 22383 (University of Illinois Urbana-Champaign) and Carle Foundation Hospital 18CCC1757. All the study participants had no history of prostate cancer. The age ranged from 50 to 90 years with a BMI less than 50. Patients were excluded if they were being treated for sexually transmitted infection or urinary tract infection; were taking antibiotics within the past month; or were being treated for benign prostatic hyperplasia with alfuzosin (Uroxatral), doxazosin (Cardura), tamsulosin (Flomax) and terazosin (Hytrin) or abiraterone/prednisone or similar drugs. Participants were given a urine collection kit and detailed instructions on how to properly collect a ‘clean/sterile catch’. The urine collection kit contained an alcohol swab to clean the urethra and tip of the penis. Participants were directed to catch mid-stream urine in sterile vials. At least 20 mL of urine was collected from participants. The urine samples were processed immediately for culturomics work. In addition to urine, clinical data (such as diagnosis, histology, etc.) was collected for all patients on the study, per the approved HIPAA Authorization form.

### Urine culturomics

To isolate androgen producing bacteria, 100 µL urine from each patient was plated on the Blood agar, Columbia agar and Schaedler agar. These plates were incubated anaerobically at 37 °C. After 4–5 days, single colonies were picked using sterilized toothpicks and transferred to a 96-well plate containing PYG broth supplemented with 50 µM 11-deoxycortisol and 11OHAD. 11-deoxycortisol was used to identify side-chain cleavage activity (*desAB*) whereas 11OHAD as substrate was used to confirm 17β-HSDH activity where it gets converted to 11OHT when *desAB*-encoding microbes are lacking in the sample. After incubation in an anaerobic chamber for 5 days, each column (50 µL per well) of the 96-well plate was pooled into a single 1.5 mL centrifuge tube to extract the steroid. Columns positive for the steroid conversion were further processed to identify the individual positive well(s) and the microbe carrying the conversion activity.

### Whole genome sequencing

Urinary isolates were cultured anaerobically in PYG broth at 37 °C. Cells were collected during logarithmic growth by centrifugation at 4,000 rpm for 15 min at 4 °C. High-molecular-weight (HMW) DNA extraction, library preparation, and sequencing were performed at the Roy J. Carver Biotechnology Center (University of Illinois Urbana-Champaign). Cell pellets were washed and resuspended in 200 µL 1× PBS (Corning, VA). Enzymatic lysis was initiated with 5 µL Metapolyzyme (Sigma) followed by incubation at 37 °C for 2 h. Further lysis and DNA isolation were completed using the MagAttract HMW DNA Kit (Qiagen, MD). Following chloroform extraction, samples were centrifuged at 10,000 × g for 2 min and the aqueous phase containing genomic DNA was retained. DNA was purified using the MagMAX™ Plant DNA Isolation Kit (Thermo Fisher Scientific, MA) according to the manufacturer’s protocol. Elution was performed in 50 µL buffer, and DNA quantity and fragment distribution were assessed using a Qubit fluorometer (Thermo Fisher Scientific) and a Femto Pulse system (Agilent, CA), respectively. Purified HMW DNA was sheared to an average size of ∼10 kb using a Megaruptor 3. SMRTbell libraries were prepared with the SMRTBell Express Template Prep Kit 3.0 and sequenced on one SMRT Cell 8M using a PacBio Revio platform in CCS mode with a 30-hour movie time. Circular consensus sequence (CCS) reads were generated using SMRTLink v13.1 with the parameters: ccs –-min-passes 3 –-min-rq 0.99.

### Whole genome sequencing analysis

Raw sequencing reads were first assessed for quality using FastQC v0.11.8 (27). SeqKit v2.0.0 was used to calculate sequencing statistics including read number, total read length, and read length distribution (28). Genome assembly was performed with Flye v2.9 (29) using approximately 50-fold coverage and the parameter –-asm-coverage. Assembly completeness and quality metrics were evaluated with QUAST v5.0.2 (30) and BUSCO v5.5.0 (31). Functional annotation of assembled genomes was carried out using Prokka v1.14.6 (32). Circular genome visualizations were generated using the CGView Server (33). Taxonomic classification was assigned using GTDB-Tk v2.3.0 with database release R220 (34).

### Gene expression and protein purification

The gene CKIMDCCC_00407 from *P. obesi* CFH08 was synthesized in *E. coli* K12 codon usage (**Supplementary Table 1**) expression plasmid in the pET-IDT C His vector from Integrated DNA Technologies (IDT). For expression of recombinant protein rGAN53_RS08700, the coding sequence was synthesized as a double-stranded DNA gblock (IDT) and amplified using gene-specific primers (IDT) prior to cloning into pET-51b(+) expression vector (**Supplementary Table 1**). All constructs were transferred into *E. coli* BL21-CodonPlus (DE3)-RIL Competent Cells. Transformed *E. coli* cells were cultured in 10 mL LB broth containing either ampicillin (100 µg/mL) or kanamycin (50 µg/mL), depending on the antibiotic resistance gene encoded by each plasmid, and grown at 37 °C with shaking for 6 hours. Cultures were subsequently diluted into 1 liter of fresh LB supplemented with the appropriate antibiotic and incubated at 37 °C with aeration at 220 rpm for an additional 2 hours until the OD reaches 0.4.

Recombinant protein expression was initiated with IPTG (final concentration 0.1 mM), followed by incubation at 16 °C for 16 hours. Cells were collected by centrifugation (5,000 × g, 30 minutes, 4 °C) and resuspended in 30 mL of binding buffer (20 mM Tris, 150 mM NaCl, 10% glycerol, pH 7.9) containing 25 µL β-mercaptoethanol, 250 units benzonase (25–29 units per µL), and 15 mg lysozyme (100 mg/mL), then incubated on ice for 40 minutes. Cells were disrupted by three passes through a high-pressure homogenizer (French press) at 30–40 psi. The lysate was clarified by centrifugation (12,000 × g, 45 minutes, 4 °C), and recombinant proteins were purified according to their affinity tags. Strep-tagged proteins (ampicillin resistant plasmid) were purified using Strep-Tactin resin (IBA), with clarified lysate applied to a gravity-flow column and washed with binding buffer until no protein was detected using the Bio-Rad Bradford dye reagent (Bio-Rad), followed by elution with buffer containing 2.5 mM D-desthiobiotin (IBA). His-tagged proteins (kanamycin resistant plasmid) were purified using Ni-NTA resin (Qiagen), washed with binding buffer containing 5 mM imidazole to remove nonspecific proteins, and eluted using high-imidazole elution buffer. Protein purity was confirmed by SDS-PAGE with Coomassie Brilliant Blue G-250 staining. Protein concentrations were determined using the Thermo Scientific™ NanoDrop™ 2000/2000c Spectrophotometer based on the protein extinction coefficients and molecular weights (rCKIMDCCC_00407: molecular weight, 28.2 kDa; extinction coefficient, 4,470 M⁻¹ cm⁻¹; rGAN53_RS08700: molecular weight, 28.8 kDa; extinction coefficient, 15,470 M⁻¹ cm⁻¹).

### pH optimization assay

The pH optima of rCKIMDCCC_00407 was assessed across pH 5.0–10.0 using buffers containing 100 mM NaCl and the following buffering systems: 50 mM Na acetate (pH 5–5.5), 50 mM NaPi (pH 6–7.5), 50 mM Tris base (pH 8–9), and 50 mM glycine (pH 9.5–10). For reductive assays, reaction mixtures contained buffer at the indicated pH, 150 µM NADPH, and 150 µM androstenedione. Oxidative reactions were performed in the same buffer mixtures supplemented with 150 µM NADP⁺ and 150 µM testosterone. Each reaction was pipetted in a Nunc™ MicroWell™ 96-well flat-bottom plate with 500 µM rCKIMDCCC_00407 in a final volume of 200 µL. Reactions were incubated at room temperature for 10 minutes. Changes in absorbance at 340 nm were monitored using an Agilent BioTek Epoch 2 microplate spectrophotometer, and activity was calculated using the extinction coefficient for NADPH (ε₃₄₀ nm = 6,220 M⁻¹ cm⁻¹).

### Enzyme assays

For substrate specificity analysis, reactions were conducted with 150 µM of individual steroids, including androstenedione (AD), testosterone (T), 5α-androstenedione (5α-AD), dihydrotestosterone (DHT), 11β-hydroxyandrostenedione (11OHAD), 11β-hydroxytestosterone (11OHT), 11-ketoandrostenedione (11KAD), 11-ketotestosterone (11KT), and δ¹-Adrenosterone (1,4-11KAD), Androstenediol (5-AL), Dehydroepiandrosterone (DHEA). Each reaction mixture contained 150 µM NADPH or NADP⁺, 500 nM rCKIMDCCC_00407 and 10 nM rGAN53_RS08700, and phosphate-buffered saline (PBS), with a final volume of 1 mL. All reactions were performed in triplicate and incubated for 24 hours prior to termination. Steroid metabolites were extracted using two volumes of ethyl acetate followed by vortexing. The separated organic phase was dried under nitrogen gas, and the remaining steroid extract was subsequently reconstituted in 200 µL LC-MS grade methanol for analysis.

### Kinetics assay

Kinetic parameters were determined at the previously established optimal pH values. Initial rates were calculated using the extinction coefficient for NADPH (ε₃₄₀ nm = 6,220 M⁻¹ cm⁻¹). For oxidation assays, testosterone served as the variable substrate. Reactions were carried out aerobically at room temperature, and changes in absorbance at 340 nm were recorded continuously for 10 minutes. Each reaction (200 µL final volume) contained 50 mM NaPi, 100 mM NaCl (pH 6), 150 µM NADP⁺, testosterone (5–200 µM), and 500 nM rCKIMDCCC_00407. For reduction assays, androstenedione was used as the substrate under comparable conditions. Reaction mixtures (200 µL total volume) consisted of 50 mM Tris buffer with 100 mM NaCl, 150 µM NADPH, androstenedione (5–200 µM), and 500 nM rCKIMDCCC_00407. Spectrophotometric assays were used to evaluate the substrate range of recombinant enzymes rCKIMDCCC_00407 and rGAN53_RS08700 under the optimal pH conditions determined previously. For reductive reactions, assays containing rCKIMDCCC_00407 were conducted at pH 7.5 in 50 mM sodium phosphate buffer supplemented with 100 mM NaCl, whereas assays containing rGAN53_RS08700 were performed at pH 6.0 using the same buffer system. Each assay contained 150 µM NADPH and 150 µM steroid substrate. Reactions with rCKIMDCCC_00407 contained 500 nM enzyme, while reactions with rGAN53_RS08700 contained 10 nM enzyme. Substrates tested under these reductive conditions included androstenedione (AD), 5α-androstenedione (5α-AD), 11β-hydroxyandrostenedione (11OHAD), 11-ketoandrostenedione (11KAD), δ¹-Adrenosterone (1,4-11KAD), dehydroepiandrosterone (DHEA), and estrone (E1).

Oxidative reactions were performed separately using 50 mM Tris base containing 100 mM NaCl. Assays with rCKIMDCCC_00407 were performed at pH 9.0, whereas reactions containing rGAN53_RS08700 were conducted at pH 10.0. Reaction mixtures consisted of 150 µM NADP⁺ and 150 µM steroid substrate. Reactions with rCKIMDCCC_00407 contained 500 nM enzyme, whereas those with rGAN53_RS08700 contained 10 nM enzyme. The oxidative substrate panel included testosterone (T), dihydrotestosterone (DHT), 11β-hydroxytestosterone (11OHT), 11-ketotestosterone (11KT), androstenediol (5-AL), and 17β-estradiol (E2).

Steroid substrates were first dissolved in dimethyl sulfoxide (DMSO), and the final solvent concentration was adjusted to 6% (v/v) in all assays. Reactions were assembled in Nunc™ MicroWell™ 96-well flat-bottom plates with a total volume of 200 µL per well. Enzymatic reactions were initiated by adding the recombinant enzyme, after which absorbance at 340 nm was recorded continuously for 10 min using an Agilent BioTek Epoch 2 microplate reader. Under these assay conditions, product formation remained linear over the measurement period and across the enzyme concentration used.

Initial reaction velocities were calculated from the change in absorbance at 340 nm (ΔA340/min). Activities were then expressed relative to a reference substrate, with androstenedione used for reductive assays and testosterone for oxidative assays, each defined as 100% activity. Standard deviations were calculated from the initial velocity values before normalization and subsequently reported as a percentage of relative activity. Data represents the mean ± standard deviation from four independent experiments (n = 4).

### Liquid Chromatography Mass Spectrometry

LC-MS analyses were conducted at the University of Illinois Urbana-Champaign Mass Spectrometry Laboratory using a Waters Synapt G2-Si ESI mass spectrometer interfaced with a Waters Acquity UPLC system. Steroid metabolites were separated on a Waters Acquity BEH C18 column (2.1 × 50 mm, 1.7 µm) operated at a flow rate of 0.5 mL/min. Mobile phase A consisted of 95% water, 5% acetonitrile, and 0.1% formic acid, whereas mobile phase B contained 95% acetonitrile, 5% water, and 0.1% formic acid. The gradient was programmed as follows: 100% A from 0.0–0.5 min; linear transition to 30% A/70% B from 0.5–6.0 min; further decrease to 0% A/100% B from 6.0–7.0 min; hold at 100% B from 7.0–8.0 min; followed by re-equilibration to 100% A from 8.1–10.0 min. Samples were injected at a volume of 0.2 µL.

### Phylogenetic analysis

The amino acid sequence of the *Peptoniphilus* candidate rCKIMDCCC_00407 protein was used as a query for BLASTP against the NCBI non-redundant protein database, and the top 100 homologous protein sequences were downloaded in FASTA format. Sequences were imported into MEGA 11 (35), and a multiple sequence alignment was generated using the MUSCLE algorithm with default parameters (36). A maximum-likelihood phylogenetic tree was then constructed in MEGA 11 using the Jones–Taylor–Thornton (JTT) substitution model for amino acids, with uniform rates among sites and all alignment positions included in the analysis. Tree searching was performed using the nearest-neighbor-interchange (NNI) heuristic method, and branch support was assessed by bootstrap analysis with 1,000 pseudoreplicates. The resulting phylogenetic tree was visualized and annotated using iTOL for tree display and formatting(37).

## Results

### Isolation and characterization of androgen formation by *Peptoniphilus* species from human urine

Bacteriological culturing from urine collected from male subjects scheduled for prostate biopsy as well as age-matched control male volunteers (>50 years of age) was performed using strict clean catch protocols as previously reported (18). Colonies were screened for steroid metabolism in PYG broth containing 50 µM 11-deoxycortisol and 50 µM 11OHAD (**Fig 1A, B**) to test for side-chain cleavage and 17-keto reduction, respectively. Strain *P. obesi* CFH08 was isolated from a subject whose prostate biopsy was negative (**Fig. 1C**), while *P. harei* CFH36 was isolated from a subject subsequently diagnosed with prostate adenocarcinoma (stage T1C) (**Supplementary Figure 1**). After LC-MS-based identification of colonies positive for 17β-HSDH activity, which were cocci under light microscopy and SEM (**Fig. 1D**). LC-MS analysis of colony CFH08 and CFH36 revealed no conversion of 11-deoxycortisol (347.23 *m/z*) but conversion of 11OHAD (303.2 *m/z*) to 11OHT (305.2 *m/z*) co-migrating with authentic standards (**Fig. 1 and Supplementary Figure 1**). We then obtained the 1,941,268 bp draft genome sequence of an isolate that was determined to be *Peptoniphilus obesi* strain CFH08 cultured from blood agar under anaerobic conditions (**Figure 2A**). The genome was 98.60% complete (0.0% contamination) and consisted of 3 contigs (N50 = 1,307,710) assembled from 186,120 sequences ranging from 247-23,090 bp and 30.44% GC content. The genome has 1,850 predicted open reading frames, including 12 rRNA and 45 tRNA genes.

**Figure 1.**
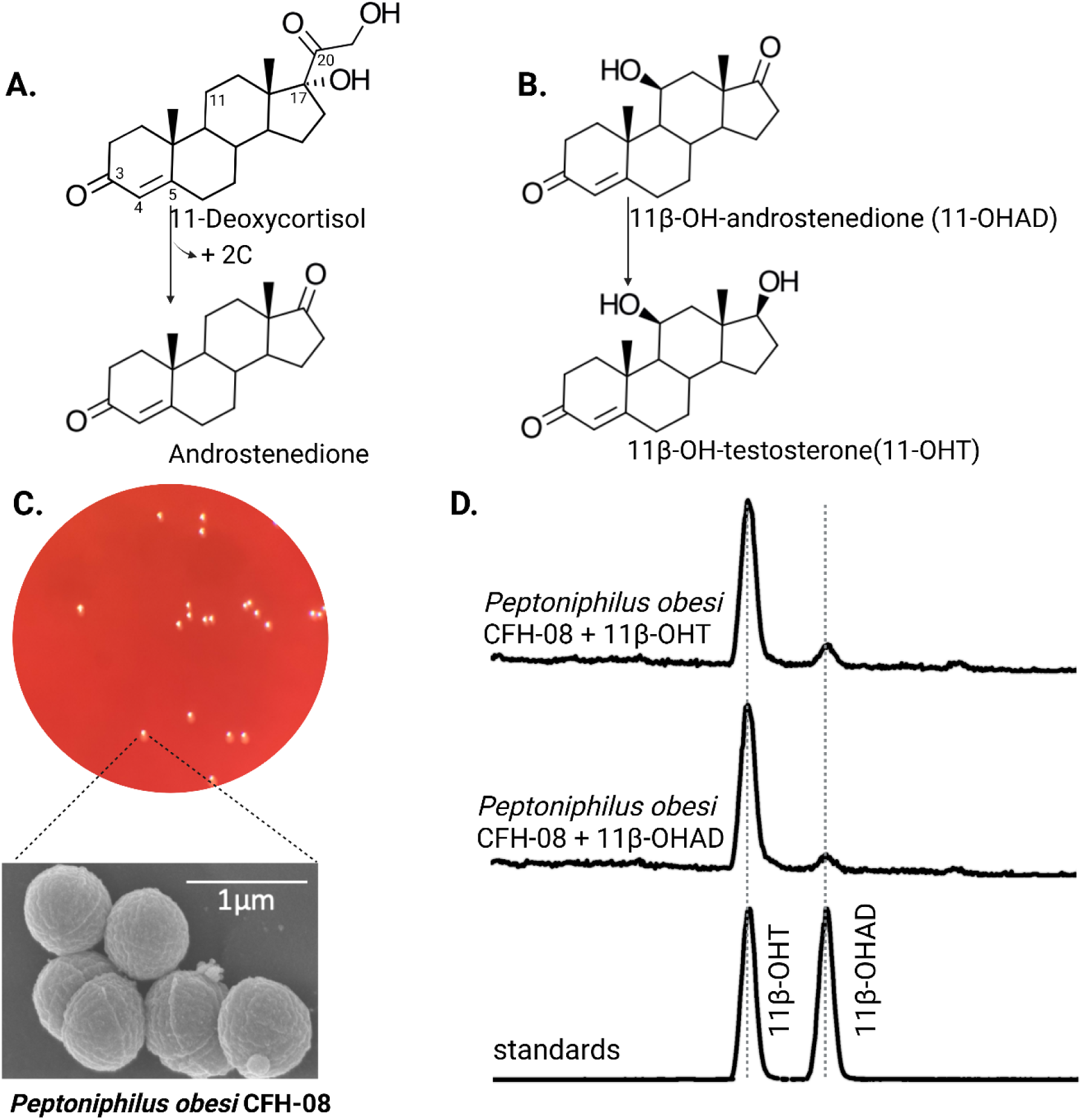
Identification of 17β-hydroxysteroid dehydrogenase activity in *Peptoniphilus obesi* isolated from human urine. A: Schematic representation of microbial side-chain cleavage of C21 glucocorticoids yielding C19 11-oxygenated steroids (+2C removal). B: Reductive 17β-hydroxysteroid dehydrogenase (17β-HSDH) conversion between 11β-hydroxyandrostenedione (11OHAD) and 11β-hydroxytestosterone (11 OHT). C: Representative colony morphology of *P. obesi* CFH08 cultured under anaerobic conditions (top) and scanning electron micrograph showing coccoid cell morphology (bottom; scale bar, 1 µm). D: LC–MS extracted ion chromatograms from whole-cell incubations of *P. obesi* CFH-08 with 11OHT or 11OHAD. Bidirectional interconversion between 11OHAD and 11OHT was observed, with product peaks co-migrating with authentic standards. Dashed lines indicate retention times corresponding to standard compounds.

**Figure 2.**
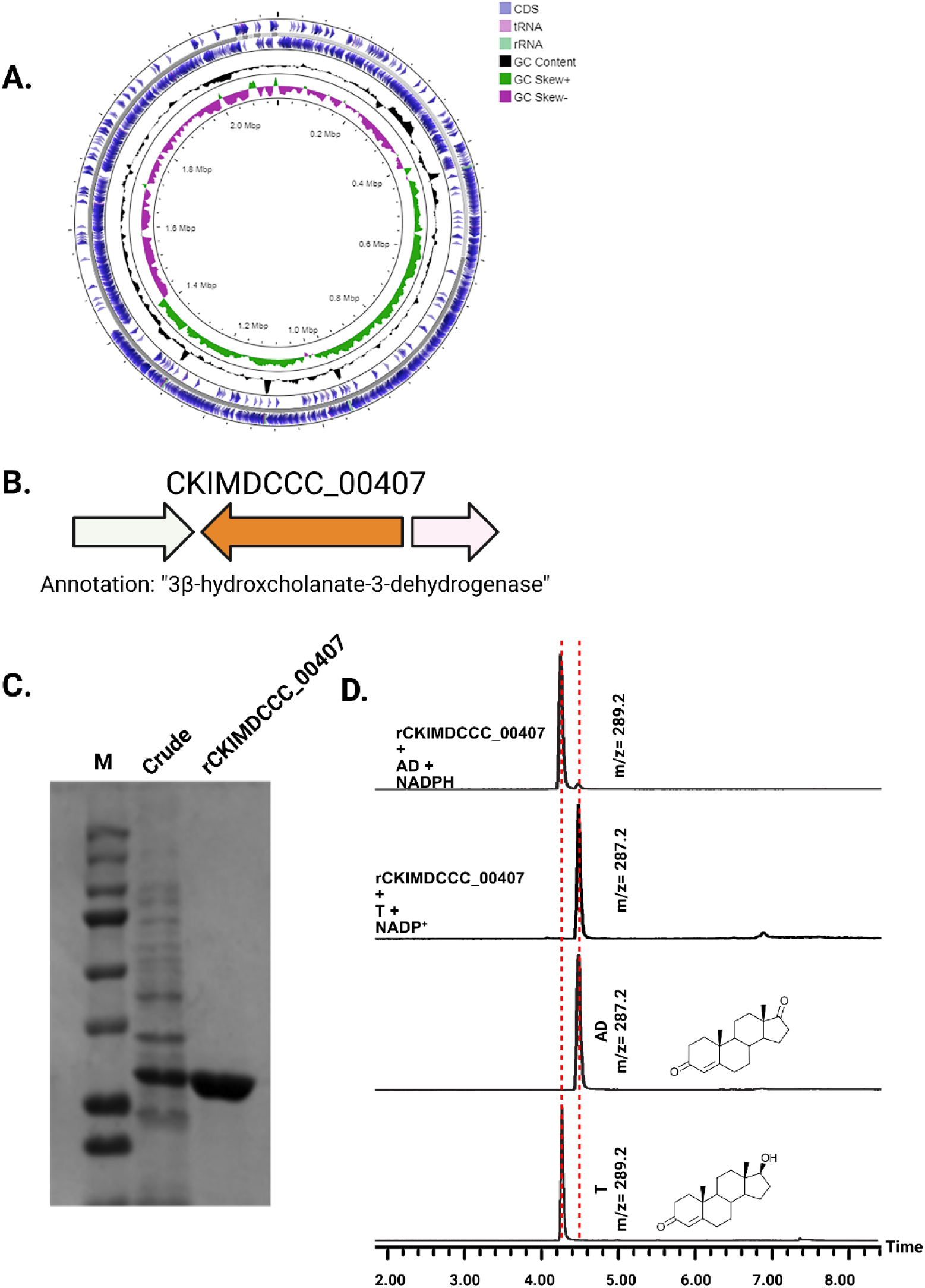
Genomic context, purification, and functional validation of rCKIMDCCC_00407. A: Circular genome map of *Peptoniphilus obesi* isolate CFH-08 showing annotated coding sequences (CDS), tRNA and rRNA genes, GC content, and GC skew (+/-). B: The gene corresponding to construct CKIMDCCC_00407 is highlighted and annotated as a putative hydroxysteroid dehydrogenase based on sequence homology. C: SDS–PAGE analysis of rCKIMDCCC_00407 during purification. Lane M, molecular weight marker; lane Crude, clarified cell lysate; lane 407, purified recombinant protein, showing enrichment of the target protein at the expected molecular weight. D: LC/MS extracted ion chromatograms demonstrating cofactor-dependent interconversion of androstenedione (AD) and testosterone (T) by rCKIMDCCC_00407. Incubation with NADPH resulted in reduction of AD to T, whereas incubation with NADP⁺ resulted in oxidation of T to AD. Authentic standards and corresponding steroid structures are shown for reference, with extracted ions indicated by m/z.

### Identification of a novel 17β-hydroxysteroid dehydrogenase

Following whole-genome sequencing of *P. obesi* strain CFH08, we sought to identify the gene(s) responsible for the observed 17β-hydroxysteroid dehydrogenase (17β-HSDH) activity. Hydroxysteroid dehydrogenases are found within the short-chain dehydrogenase/reductase (SDR), medium-chain dehydrogenase (MDR), and aldo-keto reductase (AKR) superfamilies, all of which share a conserved Rossmann fold involved in NAD(P)H binding (38). Recently, we reported the *desG* gene from the urinary tract commensal *Propionimicrobium lymphophilum*, which encodes a 26.8 kDa NADPH-dependent SDR family enzyme annotated as a “3β-hydroxycholanate dehydrogenase”.

Consistent with this annotation, genome analysis of *P. obesi* CFH08 revealed a predicted 27.1 kDa SDR family protein encoded by CKIMDCCC_00407 that was similarly annotated as a “3β-hydroxycholanate dehydrogenase” (**Figure 2B)**. Pairwise sequence alignment revealed that CKIMDCCC_00407 shares only 42% amino acid sequence identity with DesG from *P. lymphophilum* strain API-1, indicating that this enzyme represents a distinct 17β-HSDH isoform within the urinary microbiome despite catalyzing a similar reaction.

To confirm the function of CKIMDCCC_00407, the gene was synthesized and integrated into an expression plasmid (pET-IDT C-His vector) and expressed as an N-terminal affinity-tagged recombinant protein. Following heterologous expression in BL21-CodonPlus (DE3)-RIL Competent Cells and purification by affinity chromatography, rCKIMDCCC_00407 was determined to be electrophoretically pure by SDS-PAGE (**Figure 2C**). The theoretical molecular weight based on the amino acid sequence (with His-tag) is 28.2 kDa, consistent with the experimentally observed molecular weight, which was 28.2 ± 0.2 kDa.

Functional assays demonstrated that purified rCKIMDCCC_00407 catalyzed the NADPH-dependent reduction of androstenedione (AD) to testosterone (T) (**Figure 2D**). Conversely, incubation with testosterone under oxidative conditions resulted in the formation of androstenedione, confirming that rCKIMDCCC_00407 functions as a bidirectional 17β-hydroxysteroid dehydrogenase.

### Enzyme kinetics and substrate specificity for *Peptoniphilus* 17β-hydroxysteroid dehydrogenase

Next, we performed pH optimization and determined Michaelis-Menten constants and substrate specificity for rCKIMDCCC_00407. In the oxidative direction, rCKIMDCCC_00407 had a broad optimum from pH 8.0 to 9.0, with 9.0 having the highest activity (**Figure 3A**). In the reductive direction, the optimum pH was 7.5 (**Figure 3B**). For initial velocity kinetics, we utilized 0.5 µM rCKIMDCCC_00407 and tested eleven concentrations from 5 µM to 250 µM steroid while holding NADP(H) constant at 150 µM. These optimum pH conditions were utilized for kinetic and substrate-specificity analysis in each direction. In the oxidative direction, when testosterone concentration was varied as the substrate and NADP^+^ held constant at saturation as the co-substrate, the V_max_ = 77.7 ± 22.5 nmol min^-1^ mg^-1^, the *K*_M_ = 39.6 ± 3.8 µM, the *k*_cat_ = 104.5 ± 3.1 s^-1^, and the *k*_cat_/*K*_M_ = 2.6 ± 0.3 s^-1^·µM^-1^ (**Table 1**; **Figure 3C**). In the reductive direction, when androstenedione concentration was varied as the substrate and NADPH held constant at saturation as the co-substrate, the V_max_ = 56.0 ± 5.1 nmol min^-1^ mg^-1^, the *K*_M_ = 14.4 ± 1.2 µM, the *k*_cat_ = 75.9 ± 0.9 s^-1^, and the *k*_cat_/*K*_M_ = 5.3 ± 0.4 s^-1^·µM^-1^ (**Table 1**; **Figure 3D**).

**Figure 3.**
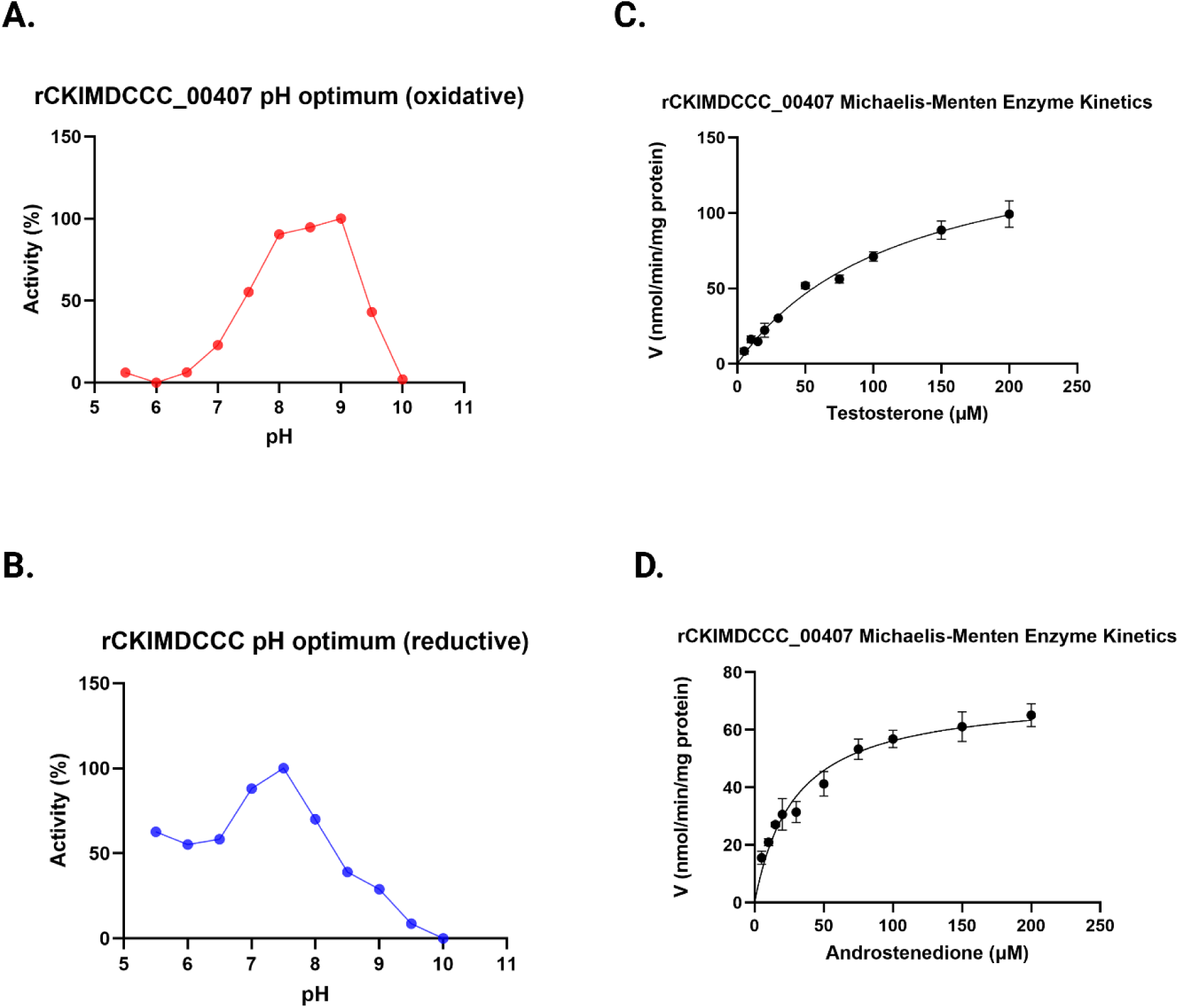
pH dependence and Michaelis–Menten kinetics of rCKIMDCCC_00407. A: Oxidative pH profile of rCKIMDCCC_00407 determined using NADP⁺ as cofactor. B: Reductive pH profile determined using NADPH as cofactor. Enzyme activity is expressed as percent relative activity across the indicated pH range. C: Michaelis–Menten kinetic analysis for testosterone in the oxidative direction. D: Michaelis–Menten kinetic analysis for androstenedione in the reductive direction. Initial reaction velocities were measured at increasing substrate concentrations and fitted to the Michaelis–Menten equation to derive kinetic parameters. Error bars represent the standard deviation of replicate measurements.

**Table 1.** Steady-state kinetic parameters of rCKIMDCCC_00407. Steady-state kinetic parameters for rCKIMDCCC_00407 determined using testosterone and androstenedione as substrates. Initial reaction velocities were fitted to the Michaelis–Menten equation to derive *Vmax* and *K_M_* values. Turnover numbers (*k_cat_*) and catalytic efficiencies (*k_cat/_K_M_*) were calculated from the fitted parameters. Values are reported as mean ± standard deviation.

| Steady state kinetics |  |  |  |  |
| --- | --- | --- | --- | --- |
| Substrate | $V_{max}$<br>(nmol min<sup>-1</sup>·mg<sup>-1</sup>) | $K_M$<br>(μM) | $k_{cat}$<br>(s<sup>-1</sup>) | $k_{cat} / K_M$<br>(s<sup>-1</sup>·μM<sup>-1</sup>) |
| Androstenedione | 56.0 ± 5.1 | 14.4 ± 1.2 | 75.9 ± 0.9 | 5.3 ± 0.4 |
| Testosterone | 77.1 ± 22.5 | 39.6 ± 3.8 | 104.5 ± 3.1 | 2.6 ± 0.3 |

Substrate-specificity analysis revealed broad substrate recognition in the reductive direction with the order of activity 11KAD > 1,4-11KAD > AD > 5ɑ-AD > 11OHAD > DHEA >> E1 (**Table 2**). The results indicate that while DHEA shares an A/B *trans* orientation with 5ɑ-AD there is nearly a 30% decrease in enzyme activity, possibly due to the 3β-hydroxyl group of DHEA. The phenolic A-ring structure of estrone (E1) resulted in the largest decrease in enzyme activity of steroids observed. In the oxidative direction, the order of preference was T > DHT > 11OHT >11KT > 5-AL > E2 (**Table 2**).

**Table 2.**
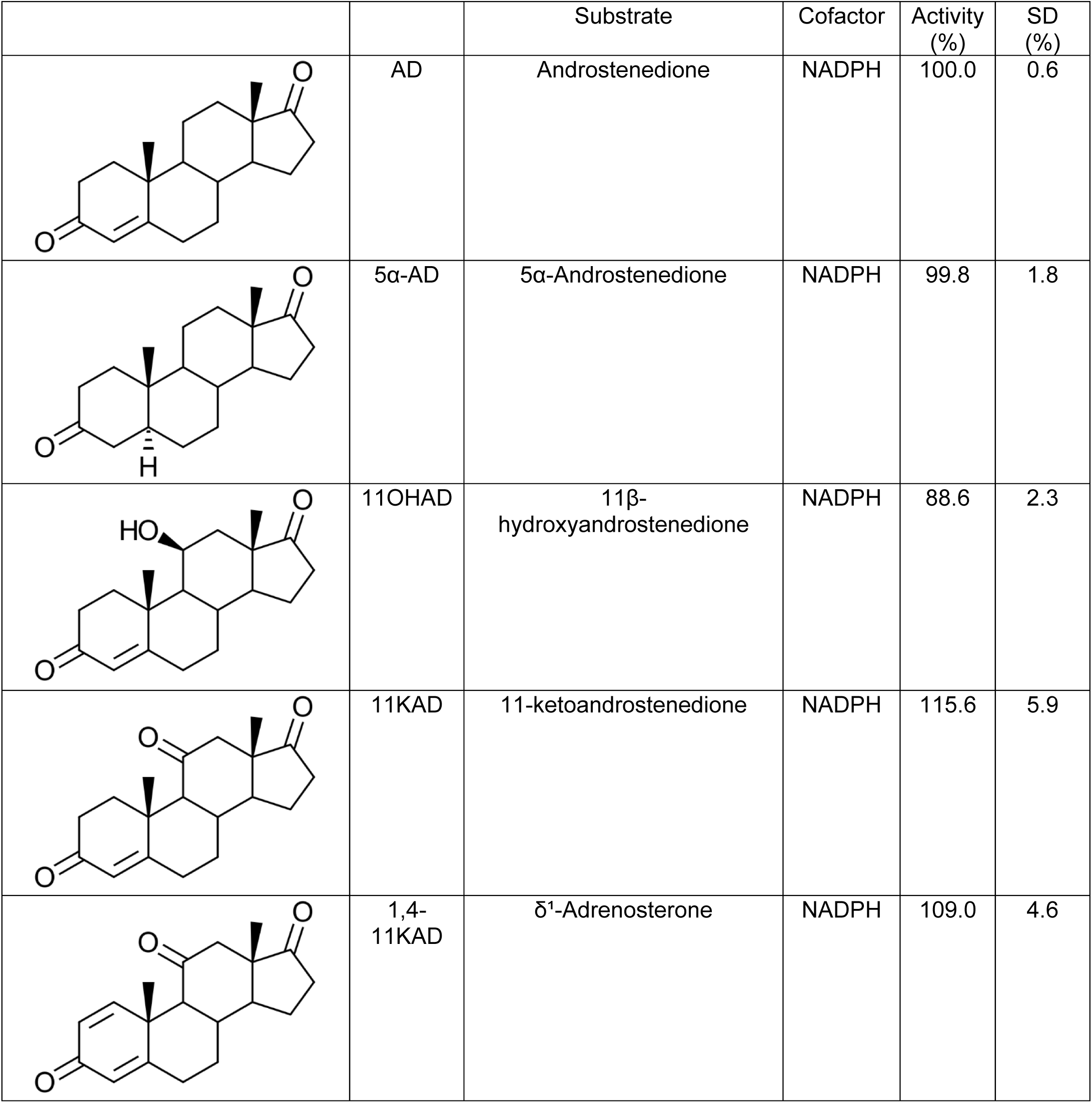
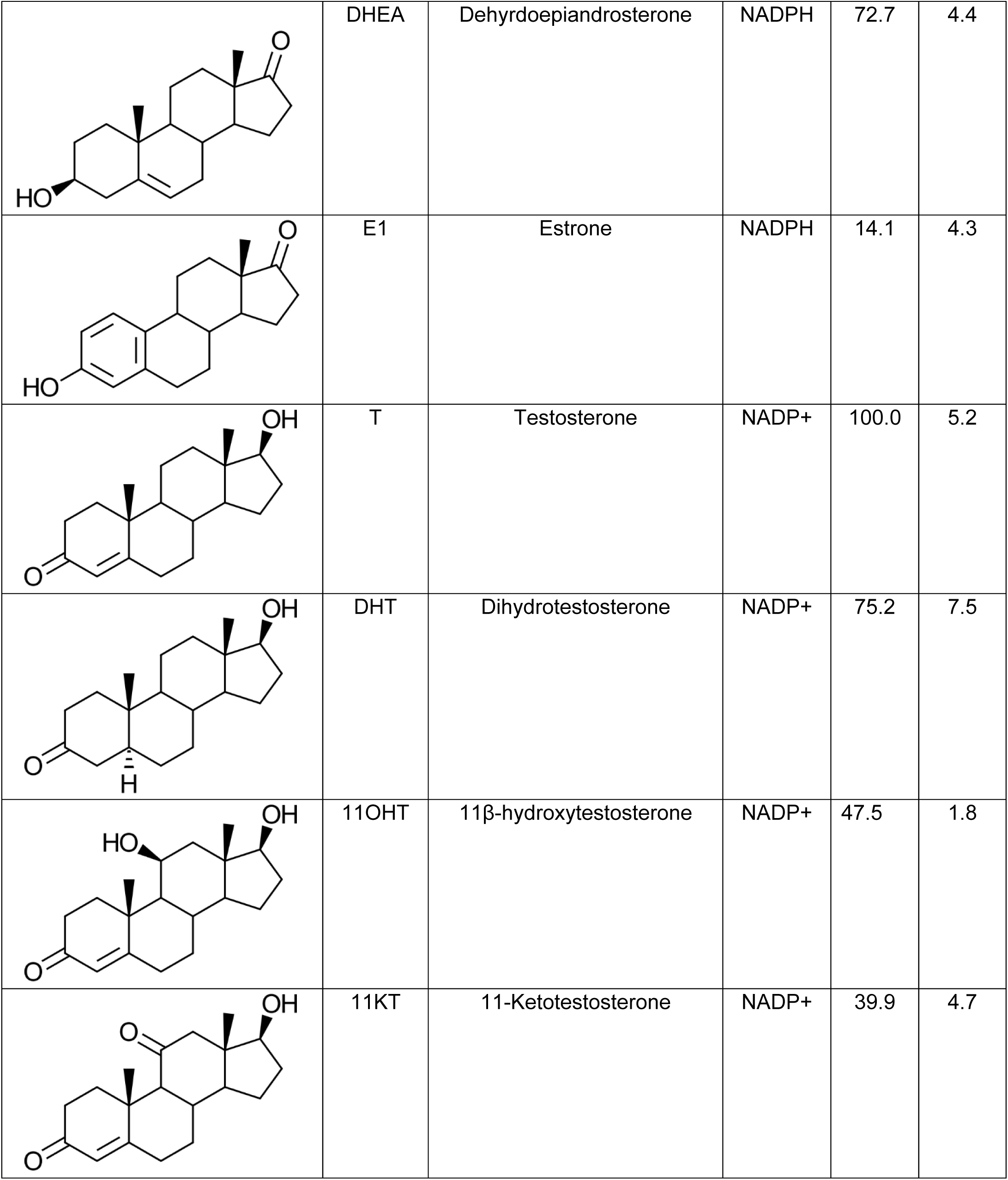
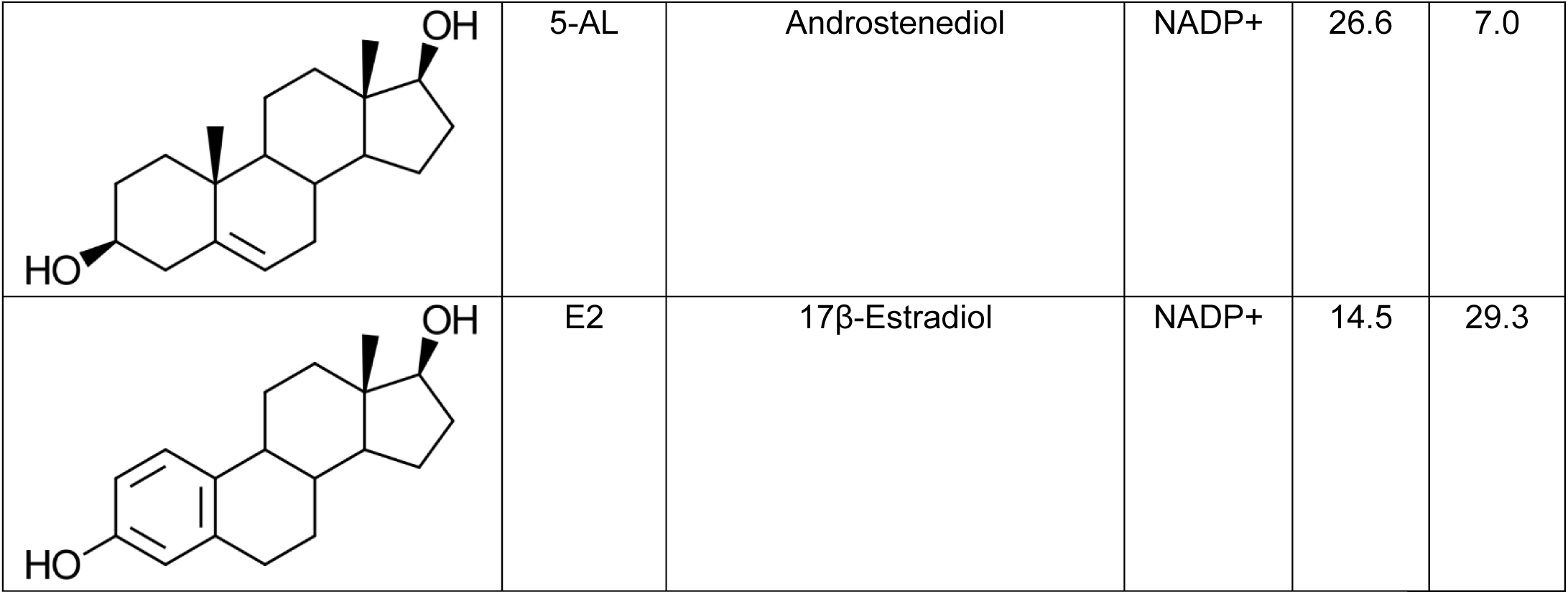
Substrate relative activity profile of rCKIMDCCC_00407. Relative activity of rCKIMDCCC_00407 toward a panel of androgen and estrogen substrates measured under identical assay conditions. Androstenedione (AD) and testosterone (T) were each defined as 100% activity within their respective cofactor conditions (NADPH or NADP⁺) and used as reference substrates to compare the activity of structurally related derivatives. Activities are reported as percentages relative to the appropriate reference substrate. Values represent mean relative activity, and standard deviation (SD) is reported as a percentage of the mean. Corresponding steroid structures are shown for reference.

### Phylogenetic analysis and characterization of a homologous 17β-HSDH from *Anaerococcus*

To explore whether homologs of the 17β-HSDH from *P. obesi* CFH08 is conserved among other host-associated microbiota, the CKIMDCCC_00407 sequence was used as a query in BLAST analyses against the NCBI protein database. The top 100 homologous sequences were selected and used to generate a multiple sequence alignment, which formed the basis for the maximum-likelihood phylogenetic tree construction (**Figure 4)**. The tree contains members of domains Bacteria and Archaea, representing several phyla including Actinomycetota, Bacteroidota, Spirochaetota, Euryarchaeota, but principally Bacillota. The query protein encoded by CKIMDCCC_00407 is within a large cluster of predominantly Bacillota sequences, interspersed with a few Actinomycetota (Coriobacteriaceae) sequences. This cluster is composed primarily of unique proteins with slight sequence variation within and between *Peptoniphilus* spp. including *P. harei, P. genitalis, P. vaginalis, P. duerdenii, P. urinale, P. urinae* (syn. *Aedoeadaptatus urinae*), and *P. obesi.* This indicates the potentially widespread expression of 17β-HSDH activity among *Peptoniphilus* spp. In addition, several other urinary tract commensals that have not previously been reported to metabolize steroids, including *Urinococcus massiliensis, Urinicoccus timonensis,* and *Actinotignum urinale*. Members of the Coriobacteriaceae, including the swine fecal isolate, *Collinsella ureilytica* (39) were also observed. Bacillota commonly isolated from the gastrointestinal tract, including *Clostridium, Ruminococcus, Coprococcus*, and methanogens including *Methanobrevibacter* were also observed in the protein phylogeny.

**Figure 4.**
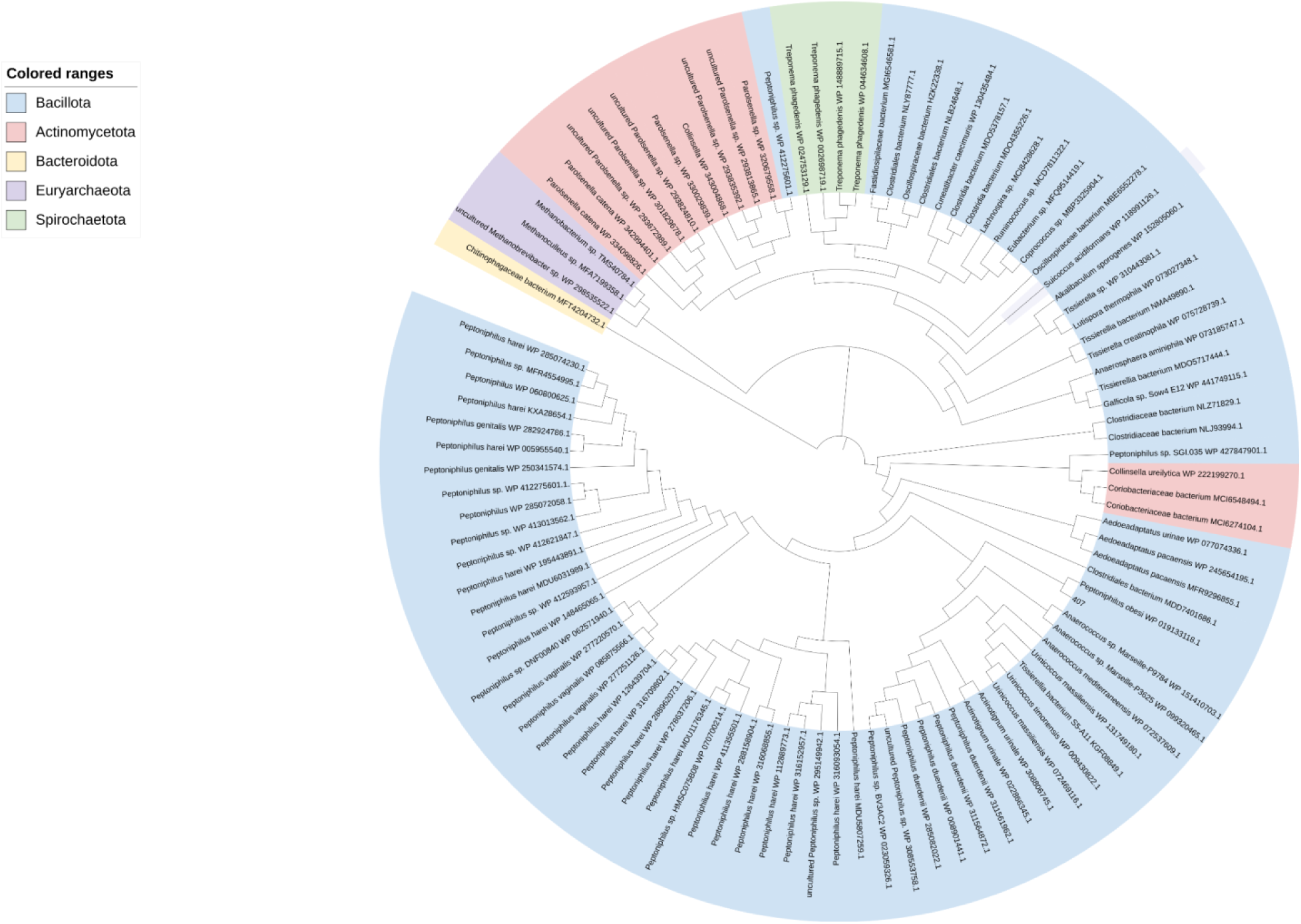
Phylogenetic analysis of hydroxysteroid dehydrogenase homologs used to identify *Anaerococcus* candidate. Maximum-likelihood phylogenetic tree constructed from amino acid sequences identified by BLAST using rCKIMDCCC_00407 as the query sequence. The top 100 homologous sequences returned by BLAST were selected for phylogenetic analysis. This approach led to the identification of a closely related homolog from *Anaerococcus*, which was subsequently selected for functional characterization as rGAN53_RS08700. Bootstrap support values are shown at branch nodes, and the scale bar represents amino acid substitutions per site.

We next wanted to “sample” from the tree to determine if closely branching neighbors also express 17β-HSDH activity. Because *Anaerococcus* has been associated in several studies with prostate cancer (20), (21), we chose to synthesize the DNA sequence corresponding to WP_151410703.1 from *Anaerococcus sp.* Marseille-P9784, annotated with the locus tag GAN53_RS08700. The sequence was codon-optimized for *E. coli* and cloned into pET-51b(+) for overexpression in BL21-CodonPlus (DE3)-RIL Competent Cells as an N-terminal streptavidin affinity-tagged recombinant protein. Following purification, protein purity was evaluated by SDS–PAGE, and enzymatic activity was confirmed by LC/MS using and AD and T as substrates **(Figure 5)**. The deduced amino acid sequence encoded by GAN53_RS08700 from *Anaerococcus sp.* Marseille-P9784 shares 86% amino acid sequence identity with the 17β-HSDH from *P. obesi* CFH-08. The theoretical *M*_r_ of rGAN53_RS08700 is 28.8 kDa while the experimentally determined *M*_r_ 29.7 ± 0.7 kDa. Hereafter, this recombinant protein will be referred to as rGAN53_RS08700.

**Figure 5.**
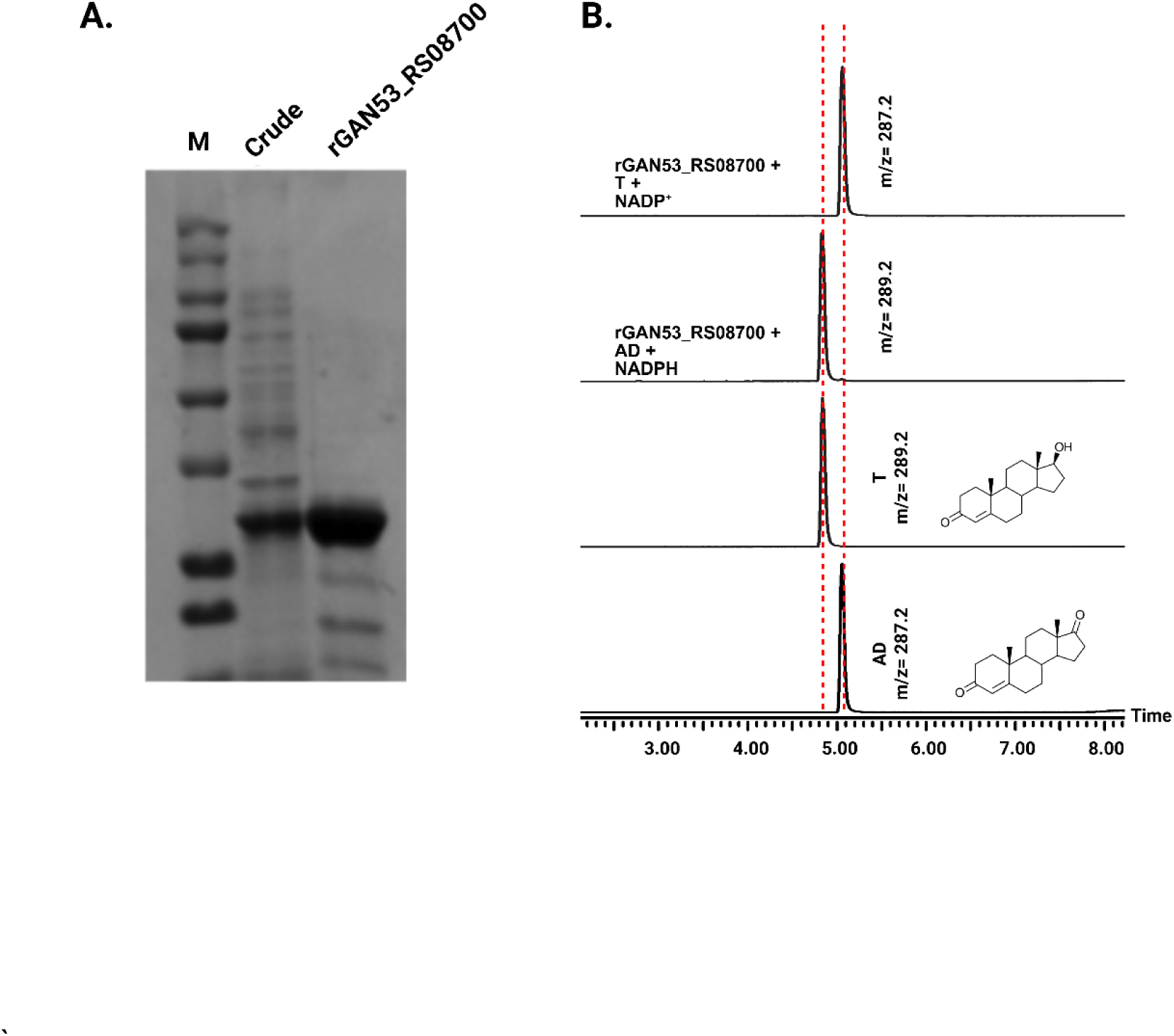
Purification and functional validation of rGAN53_RS08700. A: SDS–PAGE analysis of rGAN53_RS08700 during purification. Lane M, molecular weight marker; lane Crude, clarified cell lysate; lane GAN53_RS08700, purified recombinant protein, showing enrichment of the target band at the expected molecular weight. B: LC–MS extracted ion chromatograms demonstrating cofactor-dependent interconversion of AD and T by the purified recombinant enzyme. Incubation with NADPH results in reduction of AD to T, while incubation with NADP⁺ results in oxidation of T to AD. Authentic standards and corresponding steroid structures are shown for reference, with observed m/z values indicated.

### Enzyme kinetics and substrate specificity for *Anaerococcus sp.* 17β-hydroxysteroid dehydrogenase

Next, we performed pH optimization and kinetic analysis for the *Anaerococcus* sp. Marseille-P9784 17β-HSDH encoded by GAN53_RS08700. In the reductive diirection, rGAN53_RS08700 exhibited maximal activity observed between 6.0 and 6.5, with 6.0 having the highest activity. In the oxidative direction the maximal activity was observed between 9.5 and 10 with 10.0 having the highest activity (**Figure 6A**). These pH optima were used for all subsequent kinetics analysis. Initial velocity kinetics were measured by varying substrate concentrations from 5 µM to 300 µM while holding NADP(H) constant at 150µM. In both the oxidative and reductive directions, reaction velocities displayed sigmoidal behavior, therefore, kinetic parameters were determined using the Hill equation rather than the Michaelis–Menten model (**Figure 6. B,C**). In the oxidative direction, when testosterone concentration was varied as the substrate and NADP⁺ was held constant at saturation, rGAN53_RS08700 exhibited an apparent V_max_ of 5851.06 ± 150.97 nmol·min⁻¹·mg⁻¹, a K_0.5 of 27.4 ± 1.8 µM, and a Hill coefficient of 3.2 ± 0.3 (**Table 3**). In the reductive direction, when androstenedione concentration was varied as the substrate and NADPH was held constant at saturation, the apparent V_max_ was 4.0 ± 0.98 µmol·min⁻¹·mg⁻¹, the K_0.5_ was 23.32 ± 0.50 µM, and the Hill coefficient was 4.65 ± 0.49 (**Table 3**).

**Figure 6.**
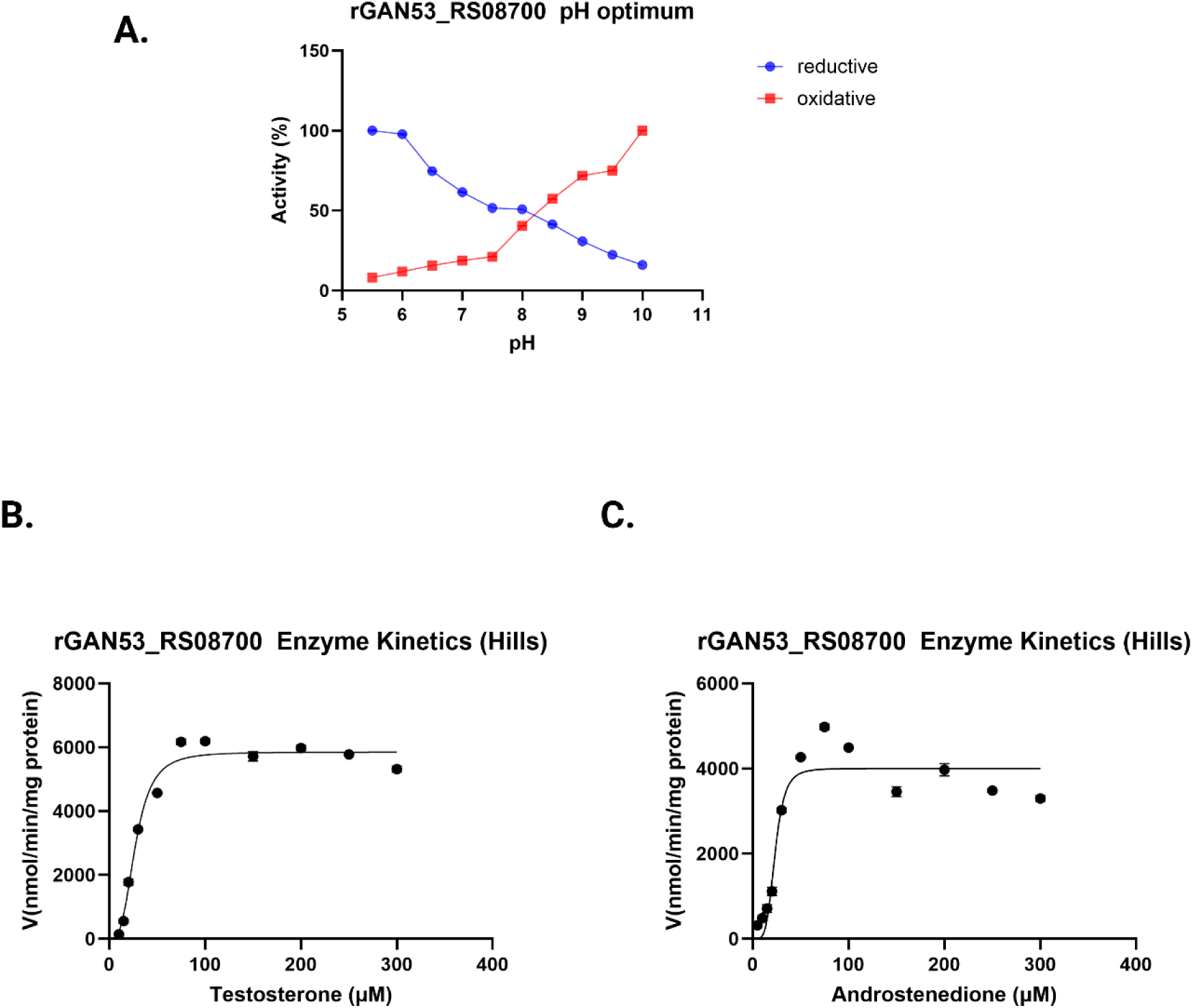
pH dependence and steady-state kinetic analysis of rGAN53_RS08700. A: pH dependence of rGAN53_RS08700 activity in the reductive (NADPH-dependent) and oxidative (NADP⁺-dependent) directions. Enzyme activity is expressed as percent relative activity across the indicated pH range. B: Hill kinetic analysis of testosterone. C: Hill kineticanalysis of androstenedione. Initial reaction velocities were measured at increasing substrate concentrations and fitted to the Hill equation to derive apparent kinetic parameters (V_0_, K₀.₅, and Hill coefficient n). Error bars represent the standard deviation of replicate measurements.

**Table 3.** Steady-state kinetic parameters of rGAN53_RS08700. Steady-state kinetic parameters for rGAN53_RS08700 were determined using testosterone and androstenedione as substrates. Initial reaction velocities were fitted to the Hill equation to derive apparent kinetic parameters (V_0_, K₀.₅, and Hill coefficient n). The Hill coefficient (n) was used to assess cooperative substrate binding. Values are reported as mean ± standard deviation.

| Steady-state Kinetics |  |  |  |
| --- | --- | --- | --- |
| Substrate | $V_0$<br>(nmol min<sup>-1</sup> mg<sup>-1</sup>) | $K_{0.5}$<br>(μM) | n<br>(Hill coefficient) |
| Androstenedione | 4002.1 ± 5.1 | 23.3 ± 0.5 | 4.65 ± 0.5 |
| Testosterone | 5851.1 ± 151.0 | 27.4 ± 0.5 | 3.2 ± 0.3 |

Substrate-specificity analysis revealed broad substrate recognition by rGAN53_RS08700 in the reductive direction, with the order of activity 1,4-11KAD > 11KAD > 11OHAD > AD > DHEA > 5α-AD >> E1 (**Table 4**). Similar to the *Peptoniphilus* enzyme, DHEA exhibited reduced activity relative to androstenedione and 11-oxygenated substrates, while estrone (E1), which contains a phenolic A-ring, resulted in the largest decrease in enzyme activity among the steroids tested.

**Table 4.**
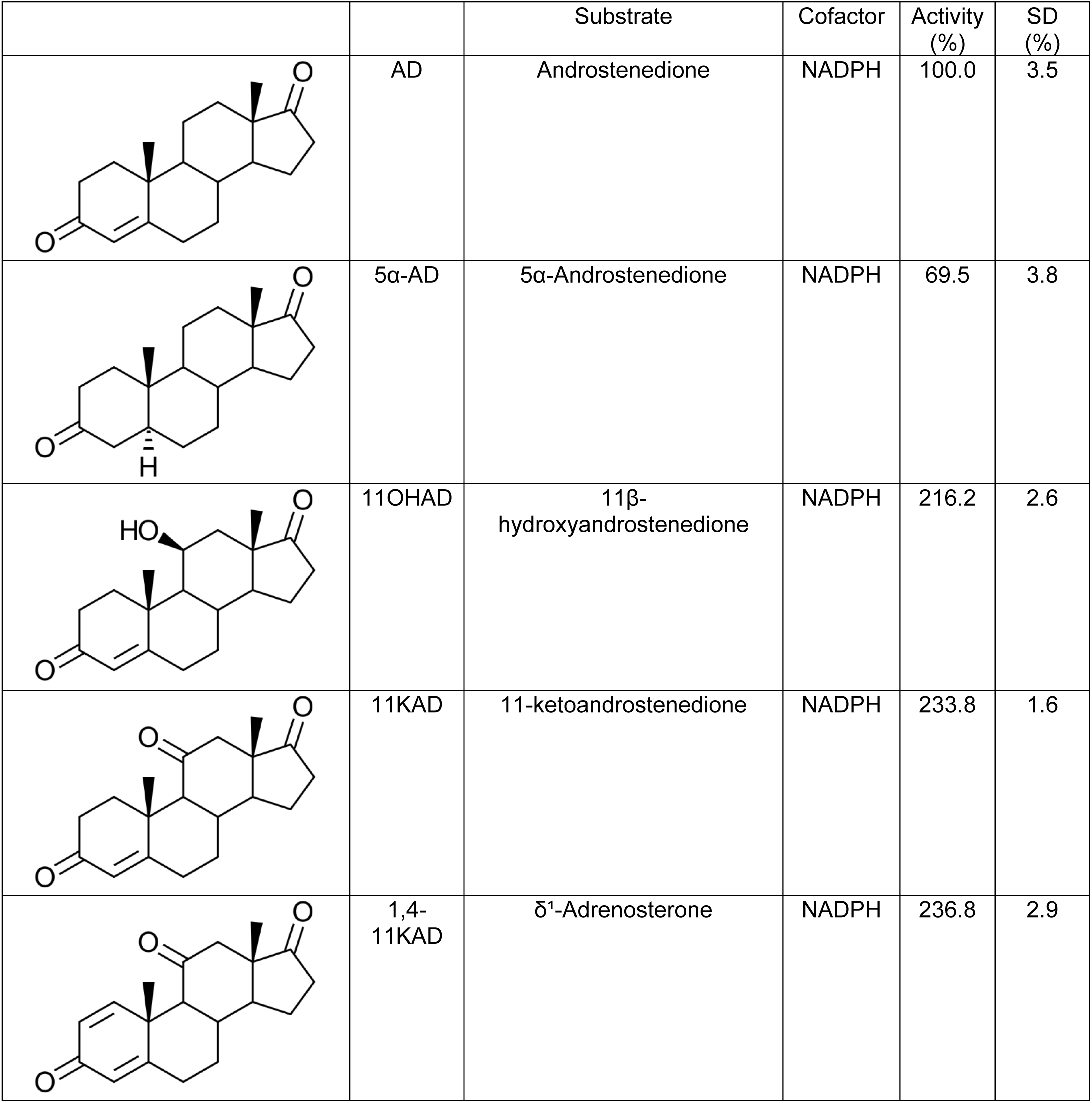
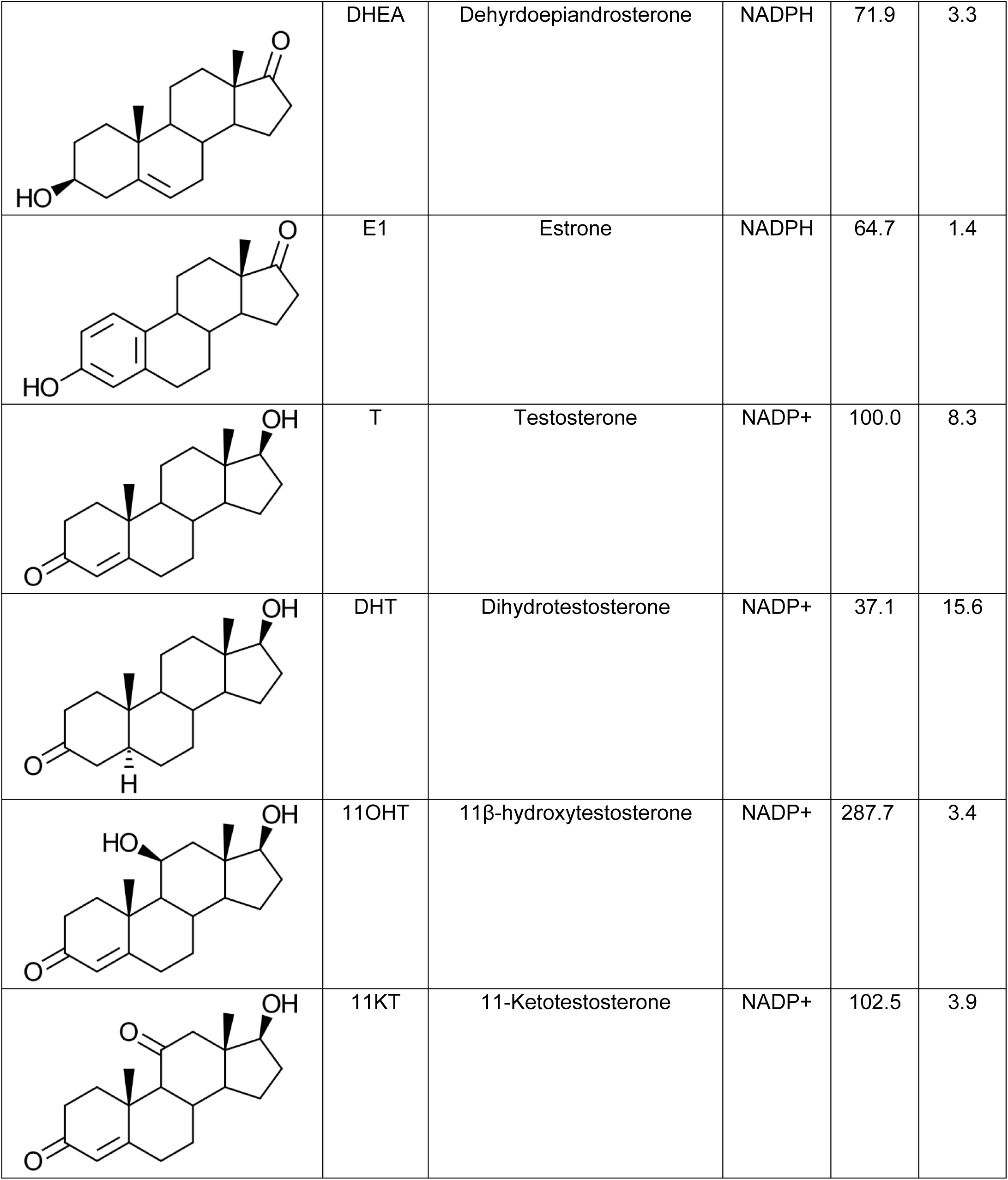
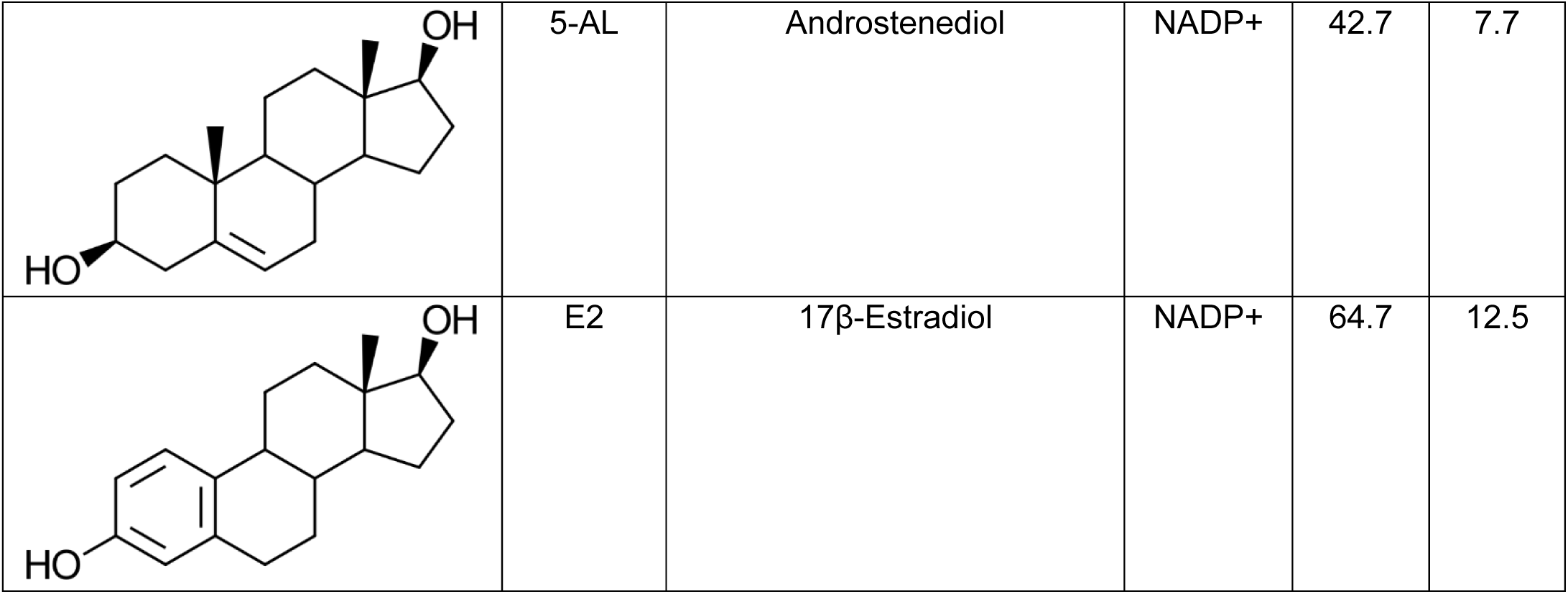
Substrate relative activity table of rGAN53_RS08700. Relative activity of rGAN53_RS08700 toward a panel of androgen and estrogen substrates measured under identical assay conditions. AD and T were each defined as 100% activity within their respective cofactor conditions (NADPH or NADP⁺) and used as reference substrates to compare the activity of structurally related derivatives. Activities are reported as percentages relative to the appropriate reference substrate. Values represent mean relative activity, and standard deviation (SD) is reported as a percentage of the mean. Corresponding steroid structures are shown for reference.

|  |  | Substrate | Cofactor | Activity (%) | SD (%) |
| --- | --- | --- | --- | --- | --- |
|  | AD | Androstenedione | NADPH | 100.0 | 3.5 |
|  | 5α-AD | 5α-Androstenedione | NADPH | 69.5 | 3.8 |
|  | 11OHAD | 11β-hydroxyandrostenedione | NADPH | 216.2 | 2.6 |
|  | 11KAD | 11-ketoandrostenedione | NADPH | 233.8 | 1.6 |
|  | 1,4-11KAD | δ¹-Adrenosterone | NADPH | 236.8 | 2.9 |
|  | DHEA | Dehyrdoepiandrosterone | NADPH | 71.9 | 3.3 |
|  | E1 | Estrone | NADPH | 64.7 | 1.4 |
|  | T | Testosterone | NADP+ | 100.0 | 8.3 |
|  | DHT | Dihydrotestosterone | NADP+ | 37.1 | 15.6 |
|  | 11OHT | 11β-hydroxytestosterone | NADP+ | 287.7 | 3.4 |
|  | 11KT | 11-Ketotestosterone | NADP+ | 102.5 | 3.9 |
|  | 5-AL | Androstenediol | NADP+ | 42.7 | 7.7 |
|  | E2 | 17β-Estradiol | NADP+ | 64.7 | 12.5 |

Notably, rGAN53_RS08700 retained measurable activity toward estrogen substrates, which was more pronounced compared to rCKIMDCCC_00407 under similar assay conditions.

In the oxidative direction, substrate preference followed the order 11T > 11KT > T > E2 > 5-AL > DHT (**Table 4**). Overall, rGAN53_RS08700 displayed a preference for both 11-keto and 11-hydroxy C19 steroids in both reaction directions, indicating enhanced activity toward these metabolites relative to non-11-oxygenated androgens. Together, these data demonstrate that rGAN53_RS08700 functions as a bidirectional 17β-HSDH with cooperative substrate binding and pH-dependent activity in both reaction directions.

## Discussion

Several independent studies have associated *Peptoniphilus* and *Anaerococcus* with prostate cancer (40, 41). Culture-based and sequencing analyses have recovered *P. harei* and *Anaerococcus prevotii* from urine, prostate tissue, and expressed prostatic secretions of men with biopsy-confirmed prostate cancer but not from controls (24, 26, 42). These taxa were also included in an anaerobic biomarker set linked to higher-risk prostate cancer groups. Despite these consistent associations, the functional activities of these organisms within the genitourinary tract have remained relatively underexplored.

In this study, we demonstrate that members of both genera encode a functional NADPH-dependent 17β-hydroxysteroid dehydrogenase capable of interconverting androstenedione and testosterone. The identification and biochemical characterization of a 17β-HSDH from *P. obesi* CFH08 expands the known repertoire of microbial enzymes involved in androgen metabolism within the urinary microbiome. Notably, this enzyme shares limited amino acid sequence identity with the previously characterized urinary tract enzyme DesG from *P. lymphophilum*.

Phylogenetic analysis further revealed that homologous 17β-HSDHs are distributed among related anaerobic taxa, including *Anaerococcus* sp. Functional characterization of the *Anaerococcus* derived homolog (rGAN53_RS08700) confirmed 17β-HSDH activity. These results, coupled with our recent reports of glucocorticoid and 17-keto-steroid metabolism (e.g. AD, DHEA) by *P. lymphophilum* and *Actinobaculum massiliense* (18, 22, 23), as well as phylogenetic analysis of urinary sterolbiome genes, suggest that androgen forming capacity is not restricted to a single lineage within the urinary microbiome, but instead may be a shared functional trait among multiple anaerobic genera enriched in urinary tract microbial communities. Moreover, homologs of *P. obesi* 17β-HSDH were also identified in taxa that inhabit the gastrointestinal tract of vertebrates. *M. smithii* in particular is interesting since it is suspected to have also acquired bile salt hydrolase (43) and bile acid 12α-hydroxysteroid dehydrogenase (44) from Bacillota through horizontal gene transfer. Further work will be needed to determine the distribution of 17β-HSDHs among the homologs reported here.

Differences in substrate-specificity between DesG and *P. obesi* 17β-HSDHs were apparent. The *P. obesi* 17β-HSDH showed a stronger preference (relative to DesG) to 11-oxygenated 17-keto steroids including the microbial side-chain cleavage products of cortisone (i.e. 11K-AD). Similar substrate preference for 11K-AD has been observed for AKR1C3 (HSD17B5) (45), an enzyme implicated in prostate cancer progression (46). Pharmacological targeting of androgenic host 17β-HSDHs is a promising approach to treating prostate cancer (47), yet with multiple isoforms of urinary microbial 17β-HSDHs which share low sequence identity (∼20-25% amino acid ID) the role of urinary tract bacteria in androgen formation should be further studied in the context of therapeutic efficacy. Indeed, recent work indicates that while abiraterone acetate is an irreversible inhibitor of host steroid-17,20-desmolase (cytochrome P450 monooxygenase CYP17A1) (48, 49), blocking conversion of pregnanolone to androgen precursors (and cortisol), bacterial steroid-17,20-desmolase (steroid transketolase) is not inhibited by abiraterone (18). Moreover, the metabolism of the abiraterone co-therapy prednisone to androgens by commensal bacteria represents a clear indication that microbial metabolism of drugs and contributions to circulating androgens should not be ignored (18). Thus, the project of elucidating the human sterolbiome (1) should be considered in lockstep with current efforts to improve androgen-deprivation therapy for hormone-sensitive castration-resistant prostate cancer (50).

The capacity of *Peptoniphilus* species to metabolize steroids may have broader implications for host endocrine signaling. *P. harei* inhabiting the GI tract have recently been shown to catalyze bile acid transformations, including conversion of 3-oxo-lithocholic acid to iso-lithocholic acid (51). Together with the androgen-transforming activity described here, these observations suggest strains of anaerobes inhabiting multiple body habitats will have distinct effects depending on the local steroid metabolome. This is particularly relevant given recent evidence that microbial bile acid metabolites can influence androgen receptor signaling pathways (52), and both secondary bile acid derivatives, including those made by *P. harei*, as well as androgens have immunomodulatory function.

Collectively, our findings indicate that commensal anaerobes of the genitourinary microbiome may contribute to local androgen availability through microbial steroid metabolism. While the current study does not establish causality between microbial androgen production and prostate cancer progression, it does identify specific microbial enzymes that could plausibly influence hormone-sensitive pathways within the prostate microenvironment. Both 17β-HSDHs characterized here recognize a broad range of 17-keto steroids, including 11-oxy-androgens and the bacterial side-chain cleavage product (1,4-11KAD) of the pharmaceutical glucocorticoid, prednisone. Future studies linking microbial enzymatic activity with host signaling pathways and clinical outcomes will be needed to determine whether microbial 17β-HSDHs represent viable targets for therapeutic or biomarkers of risk for prostate cancer and other hormone-driven diseases.

## Acknowledgments

J.M.R. would like to express gratitude to both the Cancer Center at Illinois and the Center for Advanced Study at Illinois for the financial support and protected time to pursue this work. We would like to thank Alvaro Hernandez, Director of DNA Services Facility at the Roy J. Carver Biotechnology Center at Urbana-Champaign, and Christopher J. Fields, Director of High-Performance Biology Computing at the Roy J. Carver Biotechnology Center for nucleic acid sequencing and bioinformatic assistance. We thank Furong Sun, Director of Mass Spectrometry Laboratory at Urbana-Champaign for LC-MS analysis. We thank Kristen M. Flatt for SEM images of bacterial isolates. All chemical structures were created with ChemDoodle.

This work was supported by grants from the National Institutes of Health (R01 GM145920 [J.M.R]), Cancer Center at Illinois Seed Grant [J.M.R, J.I., J.E. Jr, H.R.G.], UIUC Department of Animal Sciences Matchstick grant, and Hatch ILLU-538-916.

