## Supplementary Figures for "The Urinary Tract commensal *Peptoniphilus* spp. Encodes a Novel 17β-Hydroxysteroid Dehydrogenase"

**Supplementary Figure 1. Steroid metabolism by *Peptoniphilus* strains.** Representative LC–MS extracted ion chromatograms showing metabolism of cortisol, 11β-hydroxyandrostenedione (11OHAD), and dehydroepiandrosterone (DHEA) by *Peptoniphilus* isolates cultured from human urine. Whole-cell incubations were performed with *Peptoniphilus harei* CFH36 and *Peptoniphilus obesi* CFH08, with reactions lacking bacteria included as negative controls. Chromatograms from bacterial incubations are shown alongside authentic standards for comparison. Dashed lines indicate retention times corresponding to substrate and product peaks, demonstrating strain-specific steroid metabolism by *Peptoniphilus* species.

**Supplementary Figure 2. LC–MS analysis of steroid bioconversion by rCKIMDCCC_00407.** LC–MS chromatograms illustrating the metabolism of multiple steroid substrates by rCKIMDCCC_00407. Bioconversion reactions were performed with 11β-hydroxytestosterone (11OHT), 11β-hydroxyandrostenedione (11OHAD), 11-ketotestosterone (11KT), 11-ketoandrostenedione (11KAD), dihydrotestosterone (DHT), 5α-androstenedione (5α-AD), estrone (E1), estradiol (E2), $\delta^1$-Adrenosterone (1,4-11KAD), androstenediol (5-AL), and dehydroepiandrosterone (DHEA) using NADPH or $NADP^+$ as cofactors. Each chromatogram displays the extracted ion at the indicated m/z, with substrate and product structures shown for reference. Results demonstrate cofactor-dependent interconversion of multiple androgen and estrogen substrates.

**Supplementary Figure 3. LC–MS analysis of steroid bioconversion by rGAN53_RS08700.** LC/MS chromatograms illustrating the metabolism of multiple steroid substrates by rGAN53_RS08700. Bioconversion reactions were performed with 11β-hydroxytestosterone (11OHT), 11β-hydroxyandrostenedione (11OHAD), 11-ketotestosterone (11KT), 11-ketoandrostenedione (11KAD), dihydrotestosterone (DHT), 5α-androstenedione (5α-AD), estrone (E1), estradiol (E2), $\delta^1$-Adrenosterone (1,4-11KAD), androstenediol (5-AL), and dehydroepiandrosterone (DHEA) using NADPH or $NADP^+$ as cofactors. Each chromatogram displays the extracted ion at the indicated m/z, with substrate and product structures shown for reference. Results demonstrate that this enzyme catalyzes interconversion of multiple androgen and estrogen substrates in both oxidative and reductive directions depending on the pyridine nucleotide cofactor supplied.

**Supplementary Table 1. Synthetic DNA constructs, primers, and sequences used in this study.** This table summarizes the constructs generated for *Anaerococcus* and *Peptoniphilus*, including organism source, DNA source type, construct description, nucleotide sequences (5′–3′), and vendor. For *Anaerococcus* (rGAN53_RS08700), a synthetic gBlock DNA fragment was used as the PCR template, and gene-specific forward and reverse primers were employed to amplify the insert for downstream applications. No native genomic DNA was used for *Anaerococcus* constructs. For Peptoniphilus (rCKIMDCCC_00407), the expression plasmid was synthesized directly by IDT using a pET-based C-His vector. All primers, gBlocks, and synthesized plasmids were obtained from IDT.

#### Supplementary Figure 1

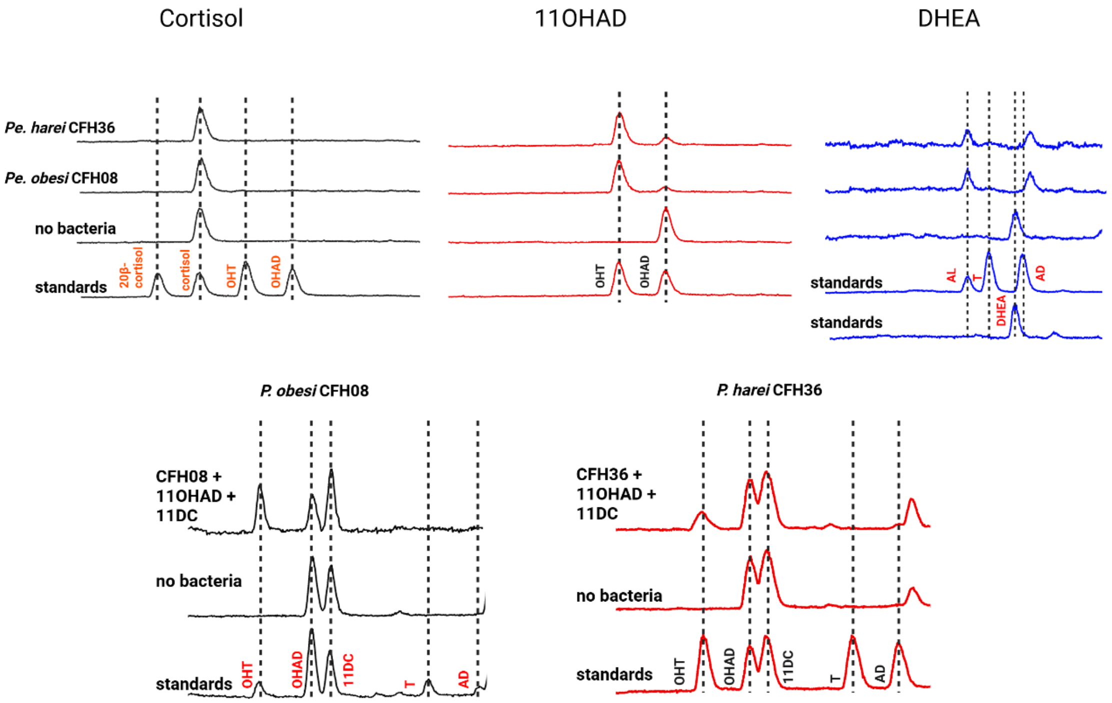

#### Supplementary Figure 2

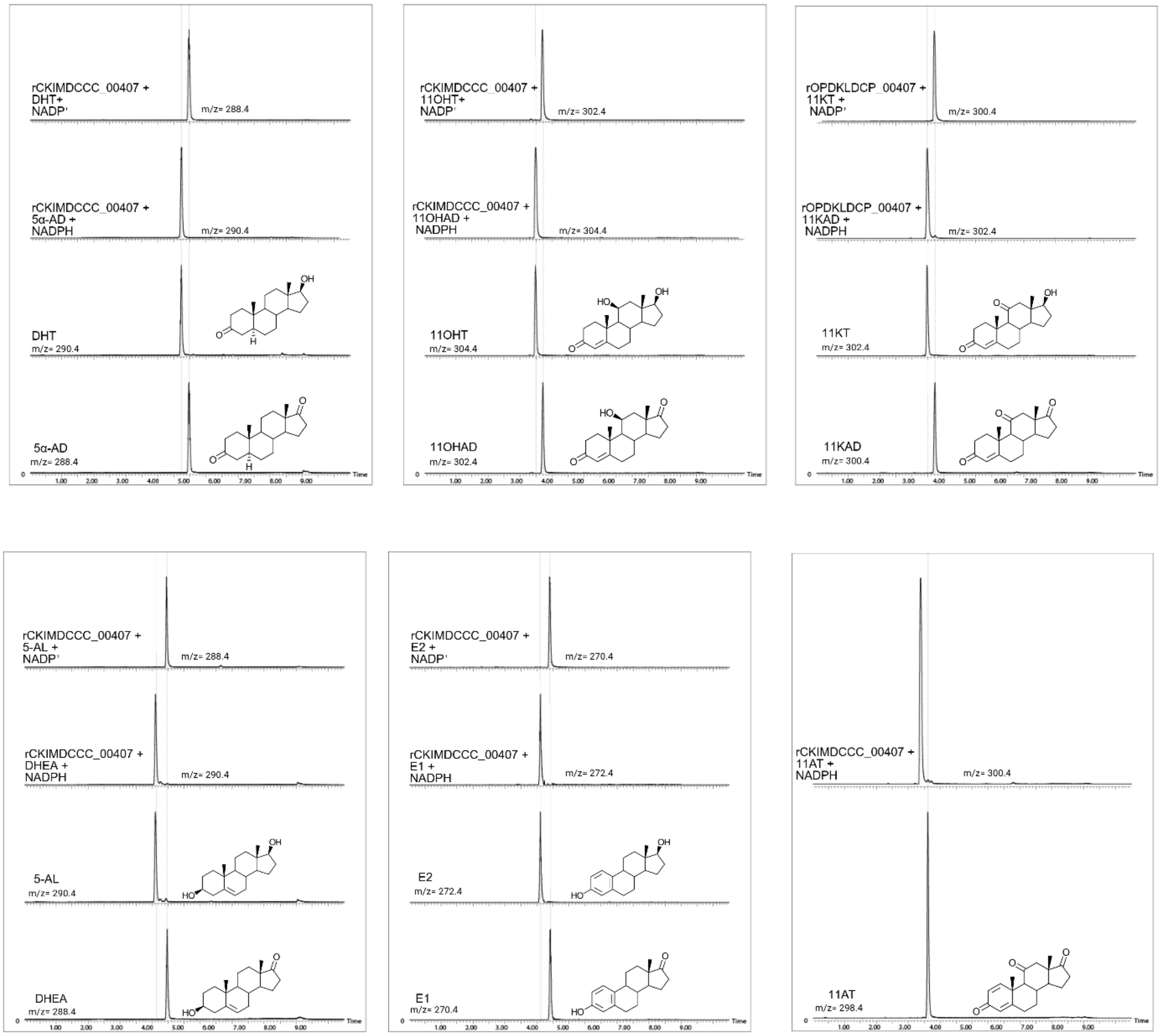

#### Supplementary Figure 3

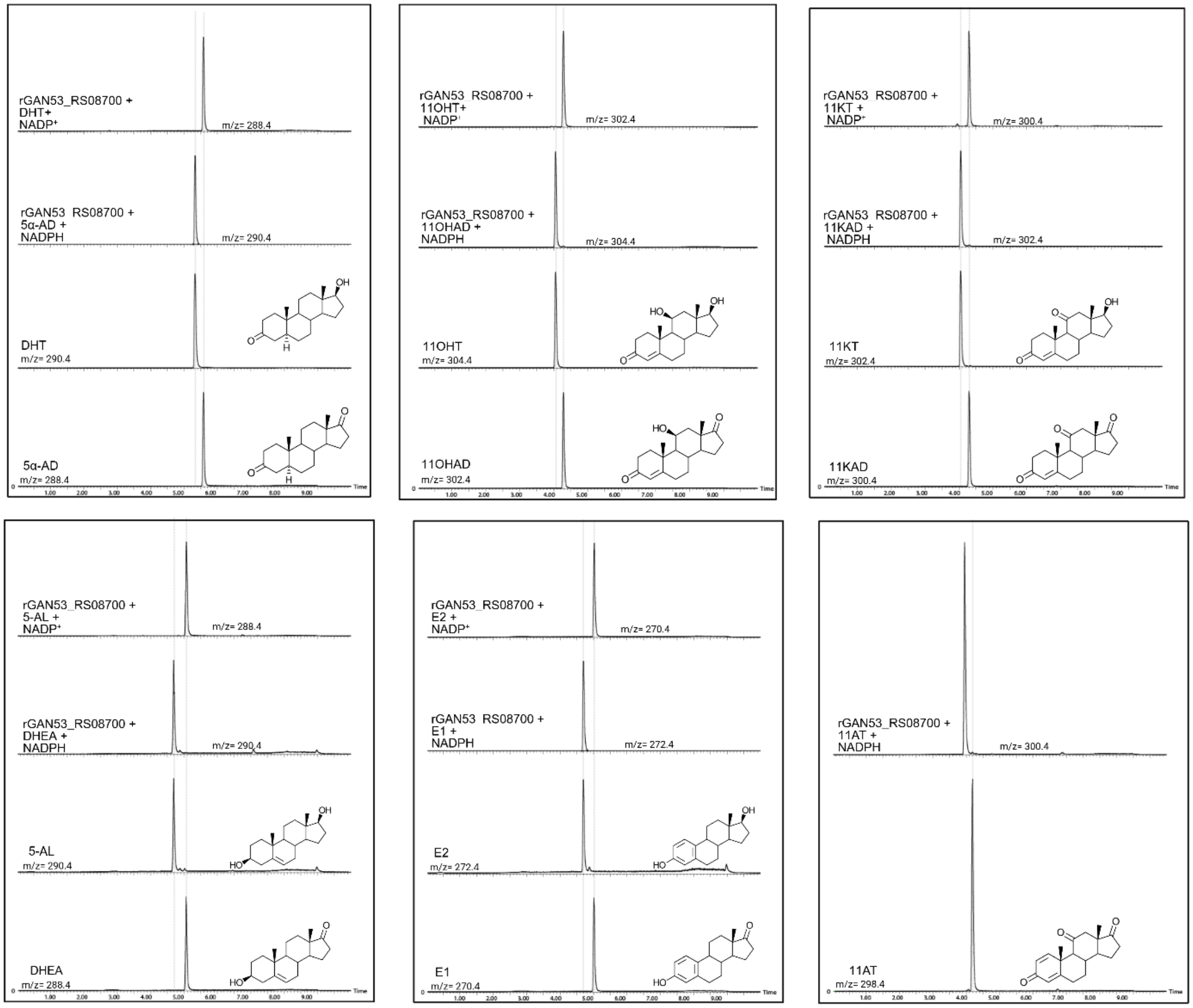

### Supplementary Table 1.

| Construct / Gene ID | Organism | DNA source type | Description | Sequence (5'–3') | Vendor |
|---|---|---|---|---|---|
| Anaerococcus GAN53_RS08700 – F primer | Anaerococcus sp. | Primer | Forward primer used to amplify synthetic gBlock | ATATATGGATCCGATGCGCTTCGAGAACAAAGTCCTGATTATCACG | IDT |
| Anaerococcus GAN53_RS08700 – R primer | Anaerococcus sp. | Primer | Reverse primer used to amplify synthetic gBlock | ATATATCAAGCTTTACATTACGGTCAGCCCACCATCAGAAACCAGAATCTGG | IDT |
| Anaerococcus GAN53_RS08700 gBlock | Anaerococcus sp. | gBlock (synthetic DNA) | Linear synthetic DNA fragment used as PCR template for cloning | ATG CGC TTC GAG AAC AAA GTC CTG ATT ATC ACG GGA GCC AGC TCT GGA GTA GGG CGC AGT GCT TCG ATT ATG GCG GCC AAA GAA GGG GCA AAA GTG TAC AGC GTC GCA CGT CGC GAG GAG AAG TTG CTG GAC TTG CGC GAT GAG GTA AAG GCA CTG AAC TGT CAG GGG GAG ATC ATC CCC GTA GTT TGC GAT GTC TCG AAA AAA GAG GAT GTG GAT AAA CTT TTT GAA AAA GTA AGC AAG GAA AAC GAG AAA CTG GAC GTA TTG GTA GCA AAT GCA GGA GTA ATG GAT AAA TTC GAA CCT GTG ACC CAT TGT GAA GAA GAC ACG TTC GAT TGG ATT TTC AAT GTA AAT GTA AAG GGC AGT TTC CGT CTG TTC AAA GGT GCT ATC CCT CTT ATG AAA GAC GGA GGA GCT ATC GTA GCT ACA ACC TCG ATC GCT GGA ATC CGT GGG GGG AAG GCC GGT GTC GCA TAC ACG ATG TCA AAA AAC GCC ATT CAT GGG CTG GTC CGC AAC ACG GCT GCT ATG TAT GCC AAT GAT AAG ATT CGC TGT AAC GCT GTC GCA CCA GGA GGA ATT GCT ACC GAA ATC TTG GAG AAC TTG AAC GAC GTA GAT GAA AAG GGT ATG GAG GTT GTG ATG CGT GGA CCT AAG CTG GAC AAG ATG GTA GCC ACG GCA GAA GAA ATT GCG TCT AAC TTG TTA TTC CTT GCG AGC GAT CAG GCC AGT AAC ATT AAT GGC CAG ATT CTG GTT TCT GAT GGT GGG CTG ACC GTA ATG TAA | IDT |
| Peptoniphilus CKIMDCCC_00407 plasmid | Peptoniphilus sp. | Synthesized plasmid | Full plasmid synthesized using pet-IDT C His vector | CGC TTT GAG AAC AAG GTC GTC GTG ATC ACC GGC GCG TCT TCT GGA GTT GGA CGC CGC GCA TCC ATC ATG GCA GCT AAA GAA GGA GCT AAA GTC TAC AGC ATT GCC CGC CGT AAA GAG AAA CTG GAA AGT CTT GCT GAG GAA GTC AAG GCA CTT AAC TGT AAG GGT AAG GTG ATC CCT GTA GTC GGG GAT GTC TCA AAA CAG GAA GAT GTT GAT CGC TTG TTC GAA AAA GTG AAA AAT GAA AAT GAA AAA CTG GAT GTC TTA ATT TCC AAT GCG GGC GTA ATG GAC AAA TTT GAG CCT ATC ACT CAC TGC GAA GAA GAG ACT TTC GAT CGT TTA TTT AAC ATT AAC GTG AAG GGG AGT TTC CGT TTG TTT AAA CGC GCG ATC CCC CTT ATG AAA GAT GGA GGG GCC ATC GTG GCA ACT ACG TCC ATC GCG GGC ATT CGC GGG GGT AAG GCT GGT GTG GCC TAC ACA GTG GCC AAA AGC GCC ATC CAC GGG CTG ATC CGC AAC ACC GCT GCG ATG TAT GCC AAC GAC AAA ATT CGC TGT AAC GGC GTG GCG CCG GGC GGC ATC CAG ACC GAA ATT CTT GAG AAC TTG AAC GAC GTG GAT GAA AAG GGG ATG GAG GTA GTG ATG CGC GGA CCA AAG CTG GAT AAA ATG GTC GCT TCT GCT GAG GAG ATT GCG AGT AAT CTT TTG TTC TTG GCT TCG GAT CAG GCA TCA AAC ATC ACG GGC CAA ATT TTG GTT TCA GAC GGT GGA TTG ACG ACA ATG | IDT |
